# Task-independent acute effects of delta-9-tetrahydrocannabinol on human brain function and its relationship with cannabinoid receptor gene expression: a neuroimaging meta-regression analysis

**DOI:** 10.1101/2021.11.01.466757

**Authors:** Brandon Gunasekera, Cathy Davies, Grace Blest-Hopley, Mattia Veronese, Nick F. Ramsey, Matthijs G. Bossong, Joaquim Radua, Sagnik Bhattacharyya, CBE Consortium

**Affiliations:** Department of Psychosis Studies, Institute of Psychiatry, Psychology and Neuroscience, King’s College London, UK; Department of Neuroimaging, Centre for Neuroimaging Sciences, King’s College London, UK; Department of Information Engineering, University of Padua, Italy; Department of Neurology and Neurosurgery, UMC Utrecht Brain Center, Utrecht University, Utrecht, The Netherlands; Department of Psychiatry, UMC Utrecht Brain Center, Utrecht University, Utrecht, The Netherlands; Institut d’Investigacions Biomèdiques August Pi i Sunyer (IDIBAPS), CIBERSAM, Barcelona, Spain; Department of Clinical Neuroscience, Karolinska Institutet, Stockholm, Sweden; Department of Forensic and Neurodevelopmental Sciences, Kings’ College London, London, United Kingdom; Sackler Institute for Translational Neurodevelopment, Department of Forensic and Neurodevelopmental Sciences, Institute of Psychiatry, Psychology & Neuroscience, King’s College London, UK; Institute of Clinical Pharmacology, Goethe - University, Theodor Stern Kai 7, 60590 Frankfurt am Main, Germany; Fraunhofer Institute of Molecular Biology and Applied Ecology - Project Group Translational Medicine and Pharmacology (IME-TMP), Theodor – Stern - Kai 7, 60590 Frankfurt am Main, Germany; Clinical Psychopharmacology Unit, University College London, United Kingdom; National Addiction Centre, Institute of Psychiatry, Psychology and Neuroscience, London, UK; Addiction and Mental Health Group (AIM), Department of Psychology, University of Bath, Bath, UK; Memory and Aging Center, Department of Neurology, University of California, San Francisco, CA 94158, USA; CIBM (Centre d’Imagerie Biomédicale), Centre Hospitalier Universitaire Vaudois (CHUV) MR section, Lausanne, Switzerland; Department of Radiology, Centre Hospitalier Universitaire Vaudois (CHUV), and University of Lausanne, Lausanne, Switzerland; Departamento de Psiquiatria da Faculdade de Medicina da USP; Faculdade de Medicina de Ribeirão Preto; Pesquisador Científico do Laboratório de Neuroimagem em Psiquiatria, Hospital das Clínicas da; School of Medicine of Ribeirão Preto, University of Sao Paulo

**Author notes:** **Corresponding Author**: Professor Sagnik Bhattacharyya MBBS, MD, PhD, Department of Psychosis Studies, Box P067, Institute of Psychiatry, Psychology & Neuroscience, De Crespigny Park, London SE5 8AF, UK. **Contributions**: Conception and Design: Bhattacharyya Data acquisition: Gunasekera, Davies, Bhattacharyya, Bossong, Ramsey, CBE collaborators Statistical analysis: Gunasekera, Radua, Veronese, Blest-Hopley, (overseen by Bhattacharyya) Data interpretation: Gunasekera, Davies, Bhattacharyya Drafting of the manuscript: Gunasekera, Davies, Bhattacharyya Critical revision of the manuscript for important intellectual content: Gunasekera, Davies, Bhattacharyya, Blest-Hopley, Bossong, Ramsey, Radua, Veronese Obtained funding: Not applicable Technical support: Gunasekera, Blest-Hopley, Bhattacharyya, Radua, Veronese.

**Keywords:** cannabis, THC, tetrahydrocannabinol, PET, fMRI, neuroimaging, systematic, meta-analysis, reward, memory, attention

## Abstract

**Background:** The neurobiological mechanisms underlying the effects of delta-9-tetrahydrocannabinol (THC) remain unclear. Here, we examined the spatial acute effect of THC on human on regional brain activation or blood flow (hereafter called ‘activation signal’) in a ‘core’ network of brain regions that subserve a multitude of processes. We also investigated whether the neuromodulatory effects of THC are related to the local expression of its key molecular target, cannabinoid-type-1 (CB1R) but not type-2 (CB2R) receptor.

**Methods:** A systematic search was conducted of acute THC-challenge studies using fMRI, PET, and arterial spin labelling in accordance with established guidelines. Using pooled summary data from 372 participants, tested using a within-subject repeated measures design under experimental conditions, we investigated the effects of a single dose (6-42mg) of THC, compared to placebo, on brain signal.

**Findings:** As predicted, THC augmented the activation signal, relative to placebo, in the anterior cingulate, superior frontal cortices, middle temporal and middle and inferior occipital gyri, striatum, amygdala, thalamus, and cerebellum crus II and attenuated it in the middle temporal gyrus (spatially distinct from the cluster with THC-induced increase in activation signal), superior temporal gyrus, angular gyrus, precuneus, cuneus, inferior parietal lobule, and the cerebellum lobule IV/V. Using post-mortem gene expression data from an independent cohort from the Allen Human Brain atlas, we found a direct relationship between the magnitude of THC-induced brain signal change, indexed using pooled effect-size estimates, and CB1R gene expression, a proxy measure of CB1R protein distribution, but not CB2R expression. A dose-response relationship was observed with THC dose in certain brain regions.

**Interpretation:** These meta-analytic findings shed new light on the localisation of the effects of THC in the human brain, suggesting that THC has neuromodulatory effects in regions central to many cognitive tasks and processes, with greater effects in regions with higher levels of CB1R expression.

## 1.0 Introduction

The extract of *Cannabis sativa* contains more than 140 different phytocannabinoids(1). Delta-9-tetrahydrocannabinol (THC) is the most abundant and extensively investigated cannabinoid in human and preclinical studies. While there is growing interest in the therapeutic potential of THC(2–11), there is also considerable evidence of its psychotomimetic effects in healthy(12–17) and vulnerable people(18), as well as those with schizophrenia(19), and an association between THC content of recreational cannabis with a greater risk of onset(20,21) and relapse(22) of psychotic disorders. Thus, there is a pressing need to better understand the effects of THC on the human brain.

A substantial number of studies have investigated the effects of THC-rich cannabis or THC isolate using single photon emission tomography (SPECT)/ positron emission tomography (PET) to measure cerebral blood flow (rCBF)(23–31) at rest, and functional MRI (fMRI) to measure the blood oxygen level dependent haemodynamic signal during cognitive activation(32,33) to index brain function. However, conflicting results from these studies have not resulted in a clearer understanding as evident from two recent systematic reviews(33,34).

Further, the molecular underpinnings of the effects of THC on human brain function remain unclear. As the cannabinoid-type-1 receptor (CB1R), the main molecular target for THC is present throughout the brain(35,36), systemic administration of THC cannot selectively target receptors only in those brain regions involved in discrete cognitive tasks. Therefore, consistent with recent neuroimaging evidence that a core network of brain regions subserve a wide range of cognitive processes(37,38), it is likely that the diverse behavioural and neuroimaging effects of THC are, at least in part, mediated by effects on such a core network of brain regions. However, whether THC has neuromodulatory effects, that is, effects on regional brain activation or blood flow (hereafter, referred collectively as ‘activation signal’) that occur across diverse (as opposed to specific/unique) cognitive tasks and at rest on a common ‘core’ network of brain regions that subserve a multitude of processes, has never been tested.

Therefore, to answer these questions, here we first meta-analysed original studies that had examined the acute effects of THC, relative to placebo, on brain function in humans using PET, SPECT, fMRI, and arterial spin labelling (ASL), with a view to investigate which brain regions are modulated acutely by a single dose of THC in humans. We hypothesised that a single dose of THC will modulate the function of a distributed set of brain regions that are engaged across a range of cognitive tasks in line with previous literature(37,38). Specifically, we predicted THC effects on dorsal attention (superior parietal lobule extending to the intraparietal sulcus, middle temporal complex and frontal eye fields), frontoparietal (lateral prefrontal cortex, temporoparietal junction, inferior parietal lobule and anterior cingulate cortex) and visual (striate and extrastriate cortex) networks as well as on the amygdala, striatum, thalamus and lateral cerebellum. Next, we used gene expression data from the Allen Human Brain atlas(39,40), to investigate whether the effect of THC on the activation signal across different brain regions, as quantified using a meta-analytic approach, was directly associated with regional CB1R(41) and CB2R(42) gene expression. Previous studies have linked gene expression levels in the human brain with anatomical(43) and functional(44,45) indices measured using neuroimaging techniques. In accordance with current understanding about the molecular targets of THC(46) we hypothesised that the pooled estimate of the effect of THC on the activation signal across different brain regions will be directly associated with CB1R but not CB2R gene expression in these brain regions.

## 2.0 Methods

The protocol for the meta-analytic synthesis was registered in PROSPERO (CRD42019145453) and we followed recommendations for neuroimaging meta-analyses(47). A detailed description of the methods are reported in Supplementary Methods.

### 2.1 Search Strategy

A systematic search of published human literature was conducted within Ovid MEDLINE, Embase, Global Health, and PsychINFO databases in accordance with the Cochrane Handbook(48) and MOOSE guidelines(49). Search terms are detailed in Supplementary Methods.

### 2.2 Eligibility Criteria

Studies were included if they (i) assessed the effect of THC on brain function using an acute drug challenge paradigm in humans, (ii) used fMRI, PET, SPECT or arterial spin labelling (ASL) to measure brain function, (iii) conducted whole-brain analysis (thus excluding small volume correction and region of interest analyses), (iv) applied consistent statistical thresholding across brain regions, and (v) published in a peer-reviewed journal. Additional details are reported in Supplementary Methods.

### 2.3 Data Extraction

For all articles that met the inclusion criteria, authors or corresponding authors were contacted by email with a request for providing whole brain statistical maps. Some studies used multiple task contrasts, therefore, combined maps with reduced variance were calculated to avoid dependent data in the analyses(50). Where maps were unavailable, whole-brain coordinates with their t-statistic were manually extracted from the published article for the conditions of interest (THC<PLB and THC>PLB). See Supplementary Methods for further details.

### 2.4 Data analysis

Voxel-wise meta-analyses of regional brain differences were conducted using the anisotropic effect-size version of the Seed-based Mapping (AES-SDM 5.15) software package (https://www.sdmproject.com/)(51,52). For studies for which we could not obtain the map, AES-SDM uses an anisotropic non-normalized Gaussian kernel to recreate an effect-size map and an effect-size variance map for the contrast between THC and placebo from peak coordinates and effect sizes for each individual fMRI study. Once contrasts were obtained for all studies, a mean map was created by performing a voxel-wise calculation of the random-effects mean of the study maps (measured as Hedge’s g), weighted by sample size and variance of each study and between-study heterogeneity. Statistical significance was determined using standard randomisation tests(53). For details on Q_H_ statistics, Egger’s test, and jack-knife leave-one-out sensitivity analysis see Supplementary Methods.

### 2.5 Meta regression analysis: Dose

A multiple meta-regression analysis was carried out using approaches described previously(54) using a significance threshold of P< .0005(51,54). We set out to investigate the association between THC dose and pooled effect-size (Hedge’s g). To control for the confounding effect of the route of THC administration, we also entered the route of THC delivery (inhalation via respiratory tract versus oral capsule) as categorical predictor. Cook’s distance(55) was calculated to identify any studies that were a potential outlier.

### 2.6 Whole brain correlation with CNR1 and CNR2 gene expression

Detailed description of the analytic pipeline including processing of genetic data from the Allen Human Brain Atlas is reported in Supplementary Methods. In summary, from the neuroimaging data synthesis, using SDM, we extracted the effect-size estimates of the voxel of the centroid for each of the 78 regions of the Desikan-Killiany(56) atlas from our main analysis. Then, we carried out linear regression analysis with the SDM effect-size estimates for brain regions in the Desikan-Killiany(56) atlas as the dependent variable and the corresponding average CNR1 and CNR2 gene expression values derived from the Allen Human Brain Atlas as the predictor variables using Python 3.7.9(57). We followed the recommendations put forward by Arnatkevic̆iūtė and colleagues with regard to processing mRNA microarray expression data from the Allen Human Brain Atlas(39) and used the package *abagen*(58) to conduct a reproducible workflow in processing and preparing the data.

### 2.7 Subgroup analysis

To better understand sources of heterogeneity, we conducted subgroup analysis. When three or more contrasts were available, we looked at more homogeneous groups based on type of imaging activation paradigm, as well as methodological variables that may have influenced the results focusing on fMRI based studies, those that administered THC isolate, and scanner magnetic field strength.

## 3.0 Results

### 3.1 Included Studies

A final set of 22 manuscripts met the study inclusion criteria (Table 1)(12,15,67–76,59,77,78,60–66). Of these manuscripts, 17 used fMRI(12,15,72–76,59–62,67–70), 4 PET(63–66), and 1 used arterial spin labelling(71). Figure 1 shows the PRISMA flowchart(79). Twenty-three separate contrasts, derived from 22 manuscripts, were included in the analysis due to some studies reporting multiple contrasts (see Supplementary Methods). Therefore, the final sample size of participants, including those with multiple contrasts, was 372 (372 under THC condition vs 370 under placebo condition). Our key analysis included 16 studies that administered THC isolate(12,15,72,73,75–78,59–62,67,69–71) and 6 that administered THC-rich cannabis(63–66,68,74).

**Table 1.** Studies included in meta-analysis. T=Tesla, INH=inhalation, OC= oral capsule, VPA= verbal paired associates task, MIDT= monetary incentive delay task, NA= not available, DB= double blind, PC= placebo controlled, R= randomised, WS= within subject, ‘= minute, A= alcohol, C= cannabis, D= illicit drug, T= tobacco, NAD= nicotine addiction disorder

| Author | Ro<br>ute | Mode | Paradigm | Baseline<br>condition | Scanner<br>strengt<br>h (T) | Design | Sample<br>size | Mean<br>age<br>(SD) | Time to<br>scanning | Pre-scan<br>screens | Dose | THC plasma<br>level (SD)<br>ng/ml |
| --- | --- | --- | --- | --- | --- | --- | --- | --- | --- | --- | --- | --- |
| Battistella(74) | INH | fMRI | Visuo-motor<br>tacking | Visually track<br>a target | 1.5 | DB, PC,<br>R, WS | 31 | 24.1<br>(3) | 45' | A,C,D,T | 42mg | 9.3 |
| Bhattacharyya(12) | OC | fMRI | Attentional<br>processing | Oddball vs<br>standard | 1.5 | DB, PC,<br>R, WS | 15 | 26.7<br>(5.7) | 1-2h | A,C,D | 10<br>mg | 1h= 3.9 (7.3)<br>2h=5.1 (5.6) |
| Bhattacharyya(15) | OC | fMRI | VPA | Presented<br>with pairs of<br>words- state<br>if font is the<br>same | 1.5 | DB, PC,<br>R, WS | 15 | 26.7<br>(5.7) | 1-2h | A,C,D | 10<br>mg | 1h= 3.9 (7.3)<br>2h=5.1 (5.6) |
| Bhattacharyya(72) | OC | [11C]MeP<br>PEP PET<br>& fMRI | Fear processing | Neutral<br>expression | 1.5 | DB, PC,<br>R, WS | 14 | 23.8<br>(4.5) | 1-2h | A,C,D,T | 10mg | NA |
| Bhattacharyya(73) | OC | fMRI | Go/No-Go | Oddball vs<br>standard | 1.5 | DB, PC,<br>R, WS | 36 | 26.0<br>(5.5) | 1-2h | A,C,D,T | 10mg | 1h= 3.9 (7.3)<br>2h=5.1 (5.6) |
| Bossong(69) | INH | fMRI | Sternberg Item<br>Recognition | Load 1 of<br>memory<br>paradigm | 3 | DB, PC,<br>R, WS | 13 | 21.6<br>(2.1) | 5' | A,C,D,T | 6mg | 70 (40.6) |
| *Bossong(70)* | INH | fMRI | Happy/Fearful Face Matching | Sensorimotor control condition (geometric shape matching) | 3 | DB, PC, R, WS | 14 | 21.5 (2.5) | 5' | A,C,D,T | 6mg | 82.3 (45.9) |
| *Bossong(71)* | INH | ASL | Resting | NA | 3 | DB, PC, R, WS | 33 | 22.6 (4.3) | 5' | A,C,D,T | 6mg | 84.9 (43.5) |
| *Bossong(77)* | INH | fMRI | Associative memory | Pictural cue | 3 | DB, PC, R, WS | 13 | 21.6 (2.1) | 5' | A,C,T | 6mg | 58.1 (31.3) |
| *Bossong(78)* | INH | fMRI | Continuous performance task | Watch stimuli | 3 | DB, PC, R, WS | 20 | 22.9 (4.9) | 5' | A,C,T | 6mg | 78.4627.0 ng/ml |
| *Freeman(68)* | INH | fMRI | Musical Reward | Scrambled sound | 1.5 | DB, PC, R, WS | 16 | 26.2 (7.3) | 5' | C,D | 8mg | NA |
| *Jansma(67)* | INH | fMRI | MIDT | No monetary reward | 3 | DB, PC, R, WS | 10 | 25.6 (2.1) | 5' | A,C,T | 6mg | 82.8 HC 82.8 NAD |
| *Lee(75)* | OC | fMRI | Capsaicin induced pain | No pain | 3 | DB, PC, R, WS | 12 | 24- 34 | 3h | A, C,D,T | 15mg | 3.5h= 1-1.2 (estimated) |
| *O'Leary*(65) | INH | H2150<br>PET | Self-paced<br>counting task | NA | 1.5 | DB, PC,<br>NR, WS | 12 | 21.7<br>(1.4) | 10-15' | C | 20mg | Occasional=17<br>.6 (8.7)<br>Chronic=35.8<br>(19.7) |
| *O'Leary*(64) | INH | H2150<br>PET | Auditory<br>Attention Task | NA | 1.5 | DB, PC,<br>NR, WS | 12 | 30.5<br>(8.6) | 10-15' | C,D | 20mg | 2.6 (3.6)-37.1<br>(27.1) |
| *O'Leary*(66) | INH | H2150<br>PET | Auditory<br>Attention Task | NA | 1.5 | DB, PC,<br>NR, WS | 12 | 23.5<br>(4.3) | 10-15' | C,D | 20mg | 10.3 (2.5)-<br>107.2 (59.7) |
| *O'Leary*(63) | INH | H2150<br>PET | Auditory<br>Attention Task | NA | 1.5 | DB, PC,<br>R, WS | 5 | 26.2<br>(8) | 10-15' | C | 20mg | NA |
| *Rabinak*(62) | OC | fMRI | Emotional<br>processing task | Neutral<br>expression | 3 | DB, PC,<br>R, WS | 14 | 20.8<br>(2.6) | 2h | A,C,D | 7.5m<br>g | NA |
| *van Hell*(61) | INH | fMRI | MIDT | No monetary<br>reward | 3 | DB, PC,<br>R, WS | 11 | 21.7<br>(2.3) | 5' | A,C,T | 6mg | 60.1 (33.7) |
| *Walter*(76) | OC | fMRI | Visual DSDT | Control visual<br>cue | 3 | DB, PC,<br>R, WS | 13 | 25.5<br>(2.3) | 2h | A, C.D,T | 20mg | NA |
| Walter(76) | OC | fMRI | Nociceptive pain<br>DSDT | Different<br>pain intensity | 3 | DB, PC,<br>R, WS | 22 | 26.1<br>(2.9) | 2h | A, C.D,T | 20mg | NA |
| Walter(60) | OC | fMRI | Olfactory and<br>pain response | Different<br>gaseous<br>stimuli | 3 | DB, PC,<br>R, WS | 15 | 26.6<br>(2.9) | 2h | A,C,D,T | 20mg | NA |
| Winton-Brown(59) | OC | fMRI | Auditory and<br>visual stimulation | Independent<br>of sensory<br>load | 1.5 | DB, PC,<br>R, WS | 14 | 26.7<br>(5.7) | 1-2h | A,C,D | 10<br>mg | 1h= 3.9 (7.3)<br>2h=5.1 (5.6) |

**Figure 1.**
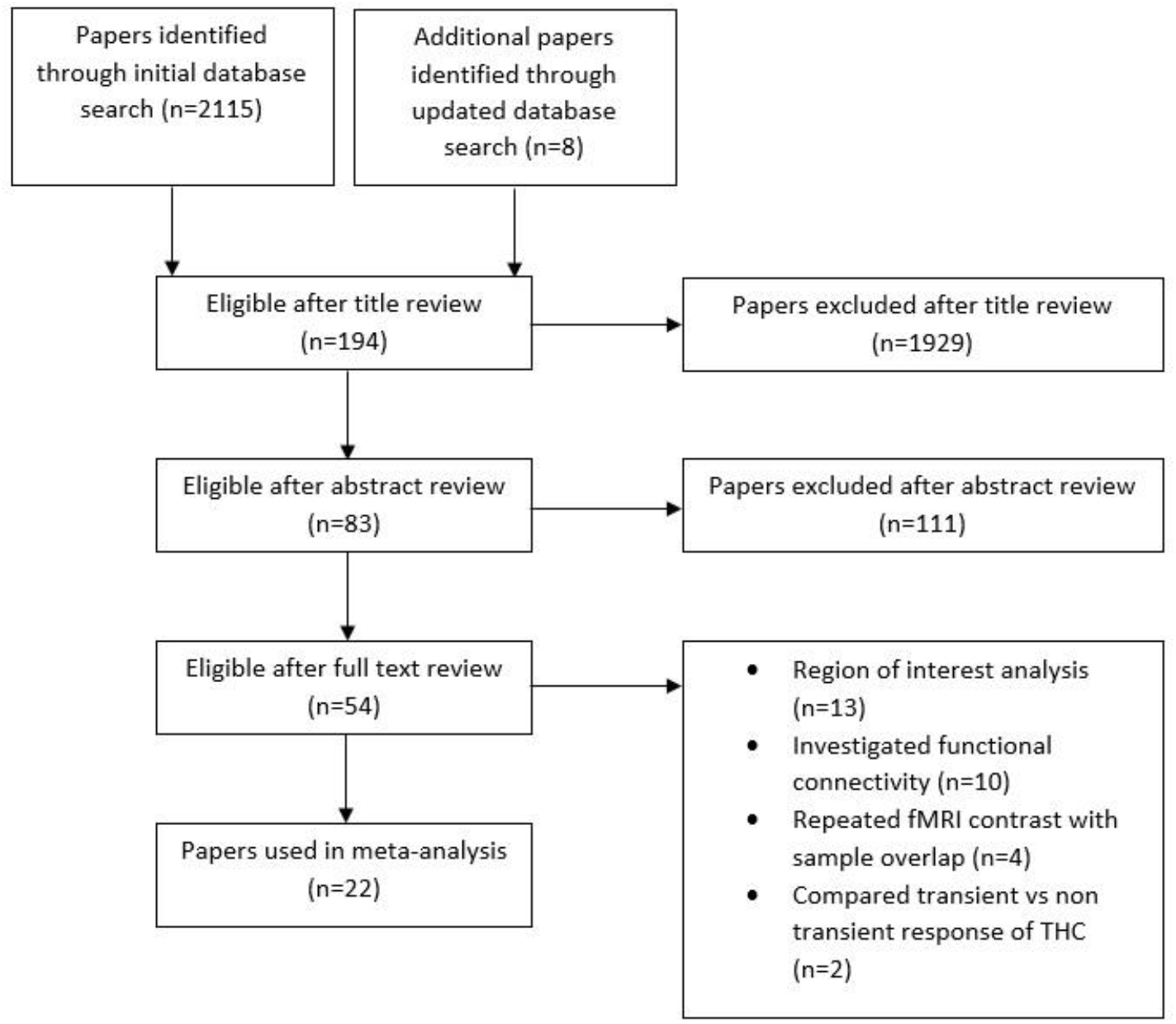
PRISMA flowchart of search strategy for meta-analysis

Studies included cognitive paradigms that engaged reward(61,67,68), memory(15,69,77), emotion(62,70,72), attentional salience(12,63,64,66,74,78) and sensory processing(59,60,75,76). One arterial spin labelling study did not use a cognitive task(71).

### 3.2 Main meta-analysis results: Effects of THC vs placebo

There were 9 regions of significantly increased activation signal (Table 2, Figure 2) under THC compared with placebo. Seven regions showed a significant attenuation of activation signal under THC compared with placebo (Table 2, Figure 2).

**Table 2.**
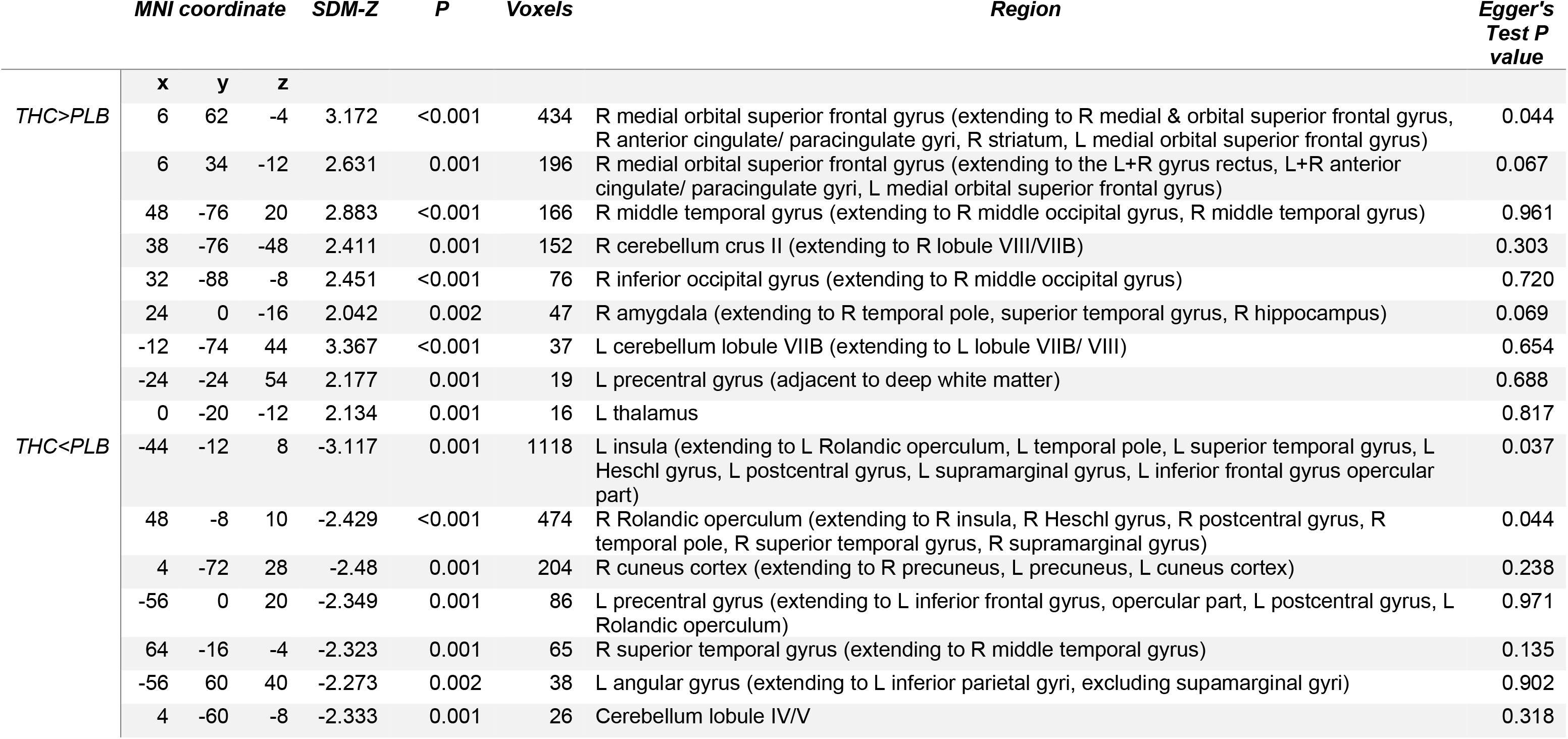
Main meta-analytic findings showing areas of increased and attenuated activation signal following THC, compared with placebo, obtained from the main multimodal meta-analysis

**Figure 2.**
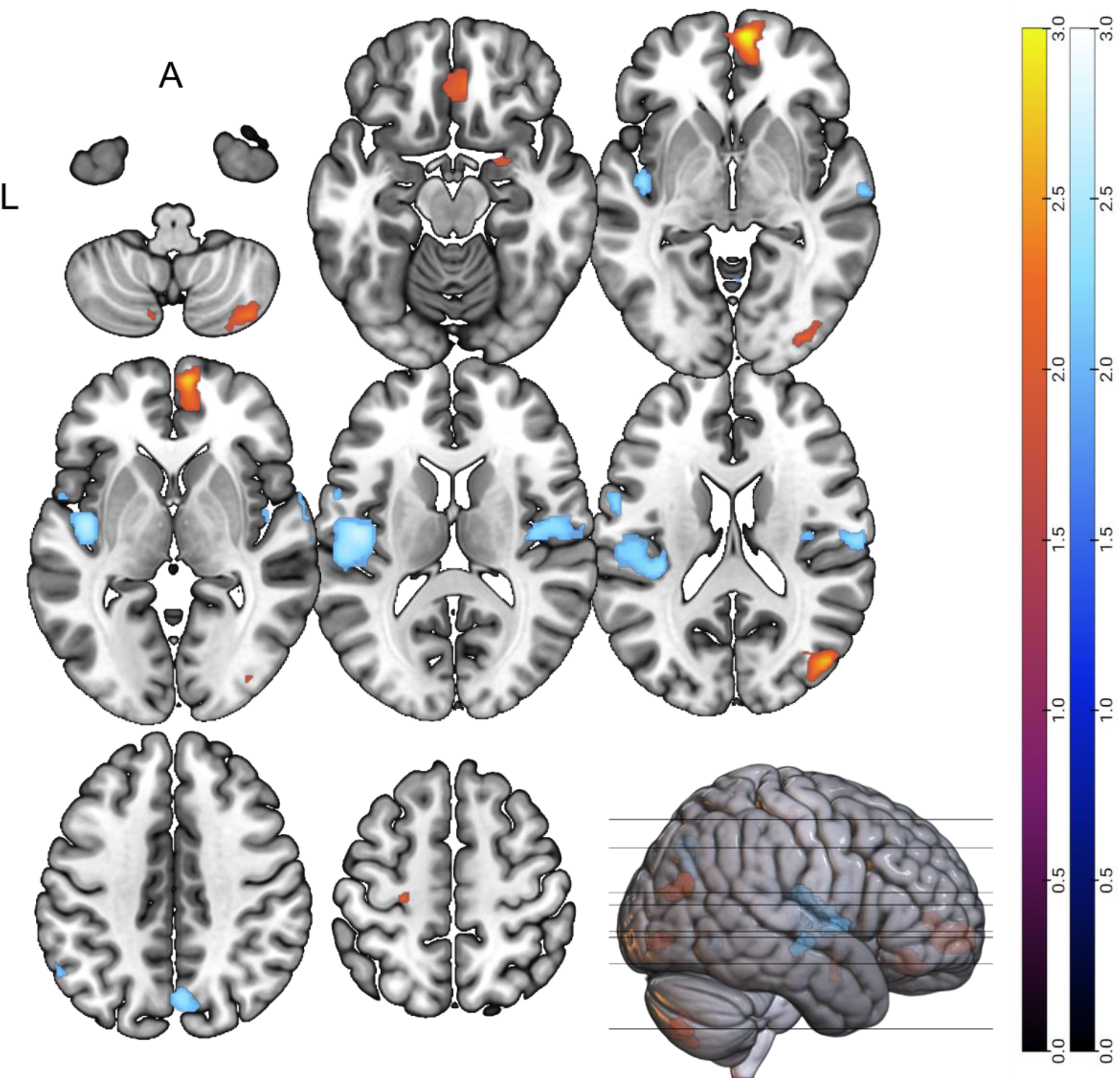
Differences in brain signal following THC compared with placebo obtained from main multimodal meta-analysis. Orange= areas of increased activation signal (THC>placebo). Blue= areas of attenuated activation signal (THC<placebo). Left side of the brain sections indicates the left side of the brain; A= anterior.

### 3.3 Sensitivity, Heterogeneity, and Publication Bias

Jack-knife sensitivity analysis showed that out of a total of 368 clusters, 87% survived following repeat analyses leaving one study out at a time (Supplementary Table 1). Funnel plots were created and examined for each cluster. Egger’s tests were performed to look for publication bias (see Table 2 and Supplementary Results). Visual inspection of overlap of meta-analytic activation maps and heterogeneity maps indicated no areas within our main analysis were significantly influenced by heterogeneity.

Different imaging modalities may be a source of heterogeneity. To ensure these factors minimally influenced our core findings, we conducted subgroup analysis of fMRI studies (Supplementary Table 7). There was significant overlap between the findings of our main results and those from the fMRI subgroup alone (Supplementary Figure 25).

Results of subgroup analyses based on cognitive paradigm and methodological variables are reported in Supplementary Tables 2-9.

### 3.4 Meta-regression analysis: Dose

Meta-regression analysis identified brain regions where there was a significant correlation between the pooled effect-size estimates of THC effect on activation signal and THC dose (6mg to 42mg) (Table 3, Figure 3).

**Table 3.**
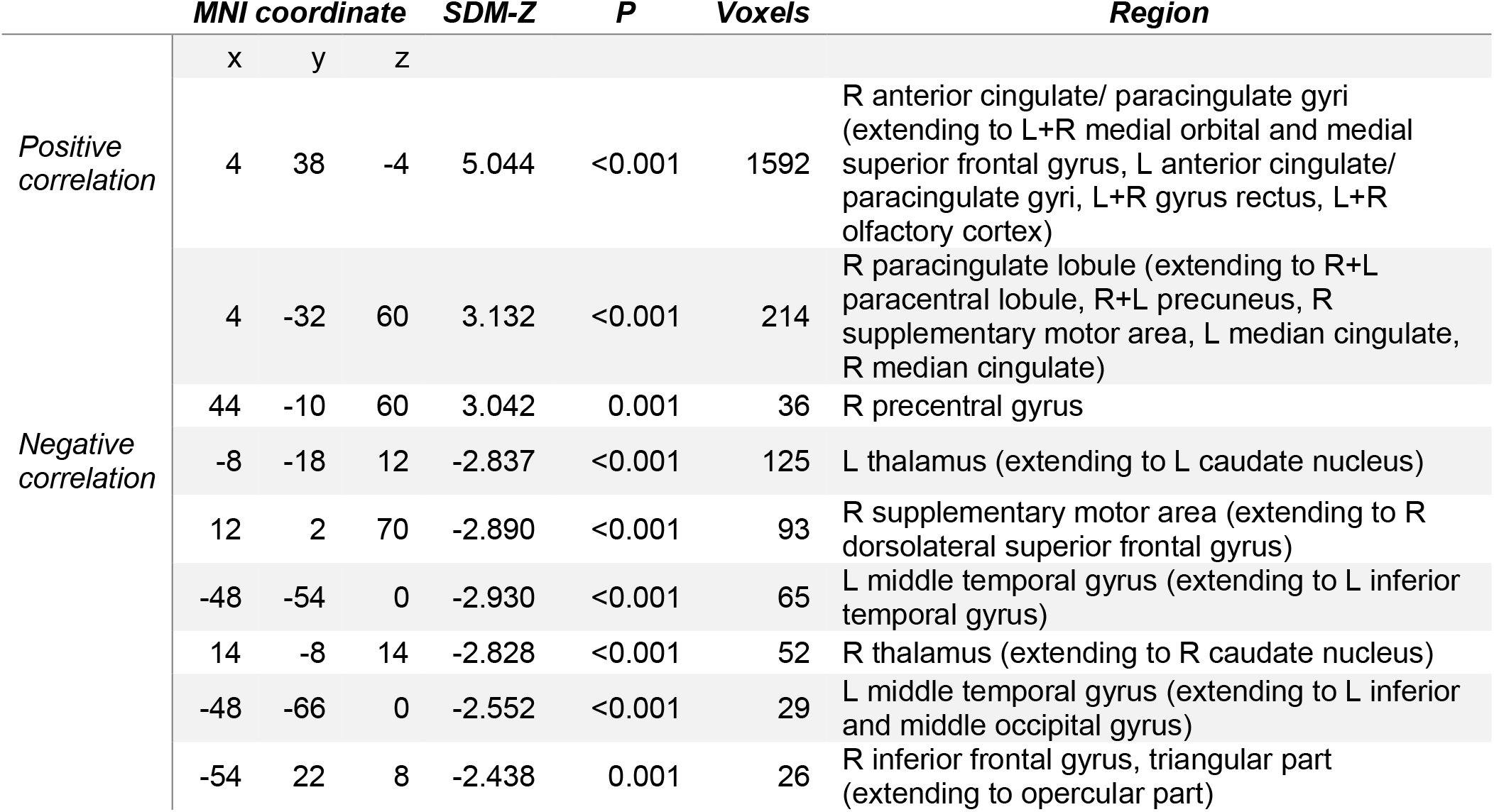
Meta-regression results showing regions where THC dose was associated with modulation of brain signal under THC compared with the placebo condition

**Figure 3.**
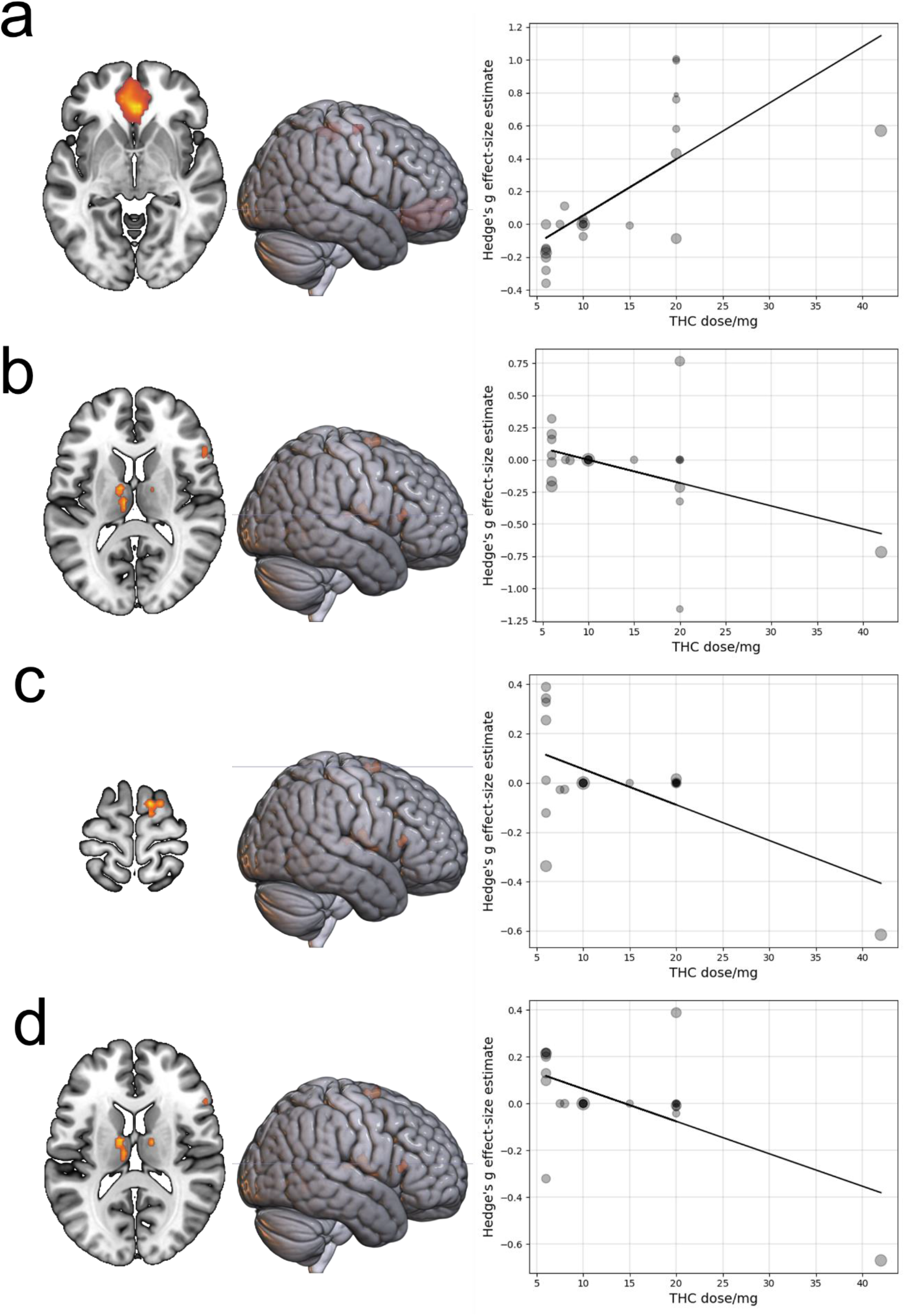
Meta-regression analysis showing relationship between THC dose (mg) and Hedge’s g effect-size estimate of brain signal modulation by THC compared to placebo. Bubble size= inverse of effect-size variance. Bubble intensity= overlap of contrasts. a) Effect-size estimates from right anterior cingulate/ paracingulate cluster b) Effect-size estimates from left thalamus cluster c) Effect-size estimates from right supplementary motor area cluster d) Effect-size estimates from right thalamus cluster

Cook’s distance(55) estimate identified the study by Battistella et al.,(74) as being a potential outlier (further discussed in Supplementary Discussion 4).

### 3.5 Whole brain correlation with CNR1 and CNR2 gene expression

Cortical and sub-cortical spatial expression of CNR1, CNR2 expression, and Hedge’s g effect size estimate of brain regions parcellated across the Desikan-Killiany(56) atlas are displayed in Figure 4. Multiple regression analysis indicated that there was a significant direct relationship between Hedge’s g effect-size estimate and CNR1 (t=2.415, P=0.018, coefficient= 0.122, 95%CI= 0.021-0.223, Figure 5) but not CNR2 gene expression (t=-0.036, P=0.971, coefficient= −0.002, 95%CI= −0.131-0.126) across the 78 brain regions of the atlas.

**Figure 4.**
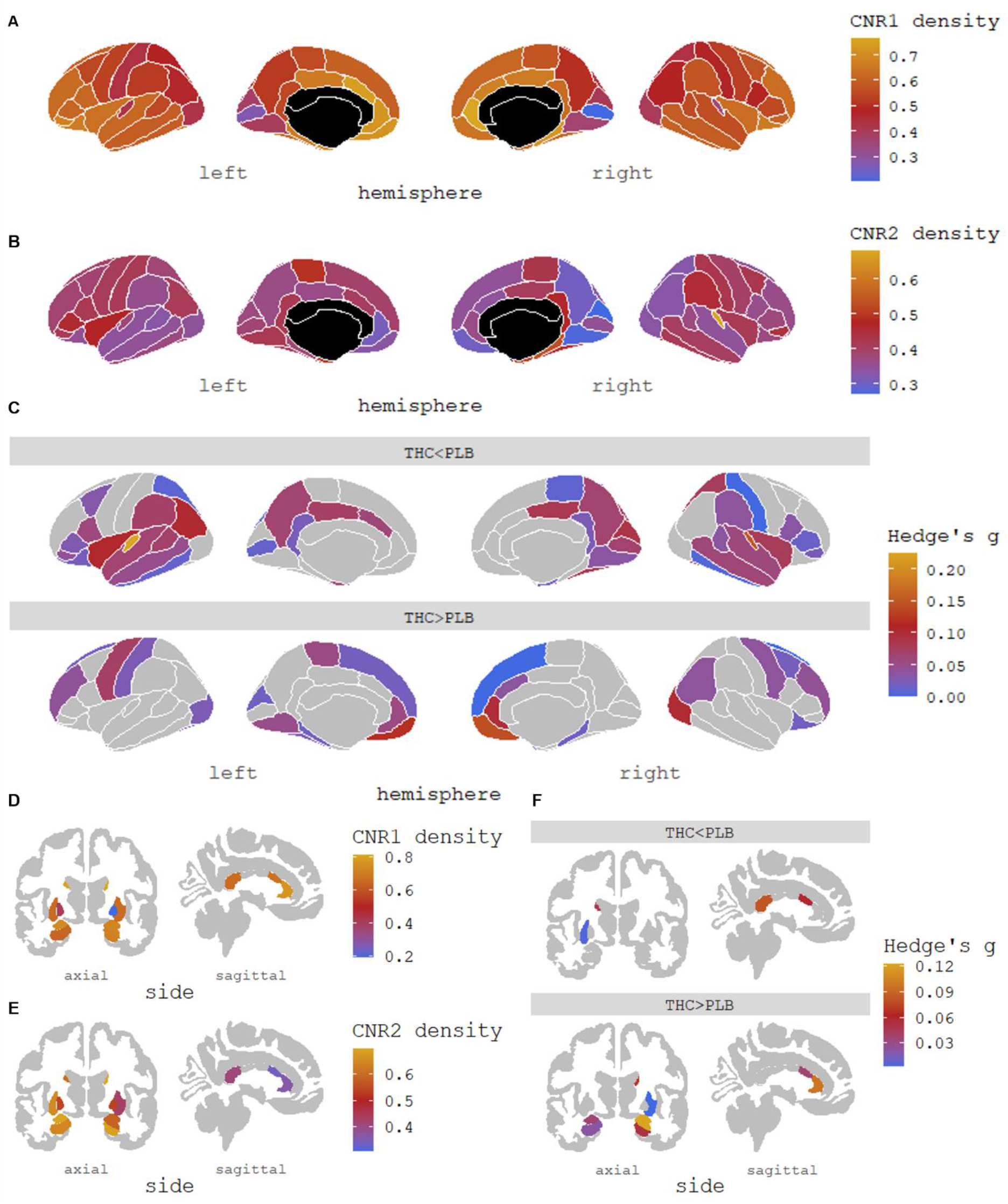
Cortical spatial gene expression of (A) CNR1 , (B) CNR2 , and (C) Hedge’s g effect size estimate derived from the main meta-analytic findings displaying regions of increased activation (THC>PLB), and attenuated activation (THC<PLB). Sub-cortical spatial distribution of (D) CNR1, (E) CNR2, and (F) Hedge’s g effect size estimate derived from the main meta-analytic findings displaying regions of increased activation (THC>PLB), and attenuated activation (THC<PLB). Figures produced using ggseg(80) in R studio(81) parcellated across 78 regions of the Desikan–Killiany brain atlas(56). Hedge’s g was extracted from the centroid of each brain parcel. Gene expression data was obtained from the Allen Human Brain Atlas(82).

**Figure 5.**
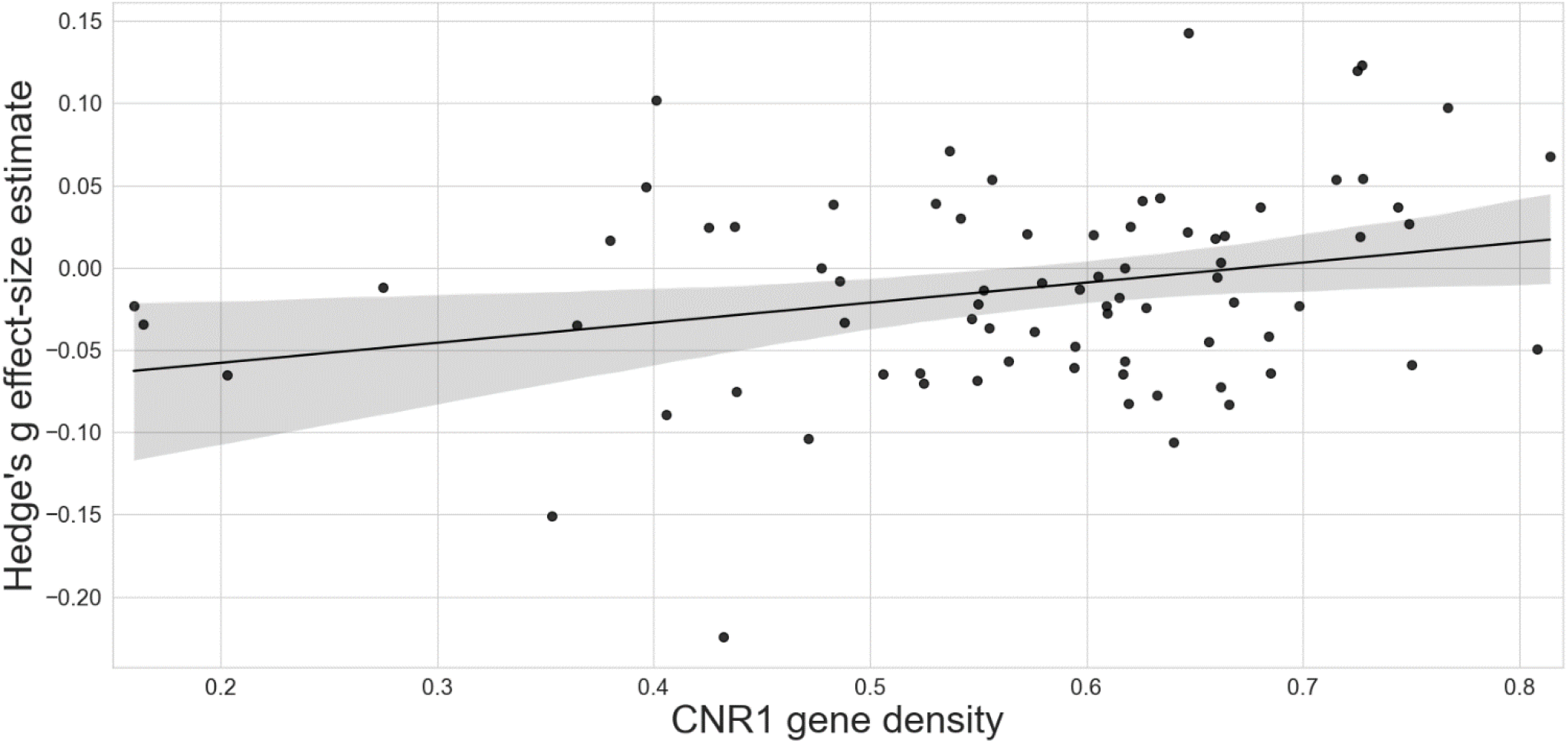
Scatterplot showing the relationship between CNR1 expression values and Hedge’s g effect size estimate of THC effect compared with placebo across the brain (based on parcellation implemented in the Desikan Killiany atlas). P=0.018, t= 2.415, R^2^= 0.073, coefficient= 0.122, 95%CI= 0.021-0.223). Shaded band around the regression line indicates 95% confidence interval.

## 4.0 Discussion

In this meta-analytic synthesis, we examined the acute effect of THC isolate and THC-rich cannabis (hereafter referred to as THC) on human brain activation signal measured using different neuroimaging modalities including fMRI(12,15,72–76,59–62,67–70), PET(63–66), and ASL(71). Using pooled summary data from 372 participants who were tested using a within-subject repeated measures design under experimental conditions acutely (5 minutes to 3 hours after administration) following a single dose of THC (ranging from 6 – 42 mg) or placebo administered orally or through inhalation, we tested whether a single dose of THC modulates the brain activation signal in a ‘core’ network of brain regions that subserve a multitude of processes. When combining data from all studies, we found that THC modulated the function of 16 brain regions. Within our predicted network of regions, THC augmented the activation signal relative to placebo in the anterior cingulate, superior frontal cortices, temporal pole, middle temporal and middle and inferior occipital gyri, striatum, amygdala, thalamus, and cerebellum crus II. There was also an attenuation of activation signal under the influence of THC in the temporal pole, middle temporal gyrus (spatially distinct from the cluster with THC-induced increase in activation signal), superior temporal gyrus, angular gyrus, precuneus, cuneus, inferior parietal lobule, and the cerebellum lobule IV/V. Further, we also found that THC augmented activation signal in regions that we had not predicted, including the paracingulate and precentral gyri (adjacent to deep white matter), gyrus rectus and the hippocampus. An attenuating effect of THC was also observed in other brain regions that we had not predicted in the insula, Rolandic operculum, Heschl’s gyrus, precentral (spatially distinct from increase in activation signal) and postcentral gyri (see Table 2 for coordinates).

Our second prediction was that the acute effect of THC on activation signal across different brain regions will be directly associated with pooled CNR1 but not CNR2 gene expression data from a set of 6 unrelated healthy volunteers (who did not take part in the neuroimaging studies reported here) in the same brain regions, as obtained from the Allen Human Brain atlas. As predicted, we found that there was a direct relationship between the effect of THC on brain activation signal with CNR1 gene expression, a proxy measure of CB1R distribution.

One of the main motivations for the present study and the analytic approach adopted here was to answer questions that previous individual studies in isolation could not address. Consistent with this objective, we identified that at the meta-analytic level, THC has effects on components of a common core network of brain regions, that has been described as a ‘domain-general’ core network that facilitates cross-task cognitive function(37). In their study, Shine et al. performed principal component analysis (PCA)(83) to identify an ‘integrative core’ network of brain regions engaged across seven diverse cognitive tasks(37) which spatially mapped onto dorsal attention, frontoparietal and visual networks as well as the striatum, thalamus, cerebellum and amygdala(37). The spatial overlap between the modulatory effects of THC that we report here and the regions within the domain-general core described by Shine and colleagues, which subserve a multitude of cognitive processes, might explain the diverse cognitive, behavioural, and neural effects of THC. Previous experimental work in cannabis users has shown that cannabis has wide-ranging effects on regional brain activation across numerous tasks(84), as well as effects on behavioural performance during those tasks(34). Please see Supplementary Discussion 1 for additional discussion regarding the effects of THC on activation signal in brain regions that were not part of the hypothesised core network, and results of analyses of cognitively homogenous subgroups of studies.

From a neurobiological perspective, effects on a common core network of brain regions makes sense: THC acts primarily via partial agonism of CB1R(36,46) which are ubiquitously distributed throughout the brain, with particularly high densities in cortex, amygdala, basal ganglia outflow tracts and cerebellum(35). THC does not selectively target CB1R only in those brain regions involved in a specific cognitive task, and instead has effects on receptors throughout the brain. In turn, THC affects the neurophysiology of these brain regions which subserve a multitude of cognitive and emotional processes. This was further demonstrated by our fMRI subgroup analysis (see Supplementary Results). We combined cognitive-specific effects from fMRI paradigms and intoxication-related effects from THC. Overlap in the brain substrates modulated by THC was observed across our main findings and the fMRI subgroup analyses. Shine and colleagues also demonstrated that the dynamic function of this integrative core is strongly influenced by the modulatory effect of neurotransmitters, and propose that any dysregulation in neurotransmitter systems, for example, in the context of neuropsychiatric disorders or as induced through pharmacological manipulation, could conceivably facilitate or impede neurotransmission through actions on this integrative core(37). In this regard, the endocannabinoid system itself may be an exemplary candidate, poised at the synapse as a critical mediator of neural homeostasis and signalling: endocannabinoids are released postsynaptically and via retrograde signalling, bind to presynaptic CB1 where they inhibit neurotransmitter release. The administration of exogenous cannabinoids such as THC may subvert this on-demand fine-tuning by indiscriminately binding CB1 receptors, and therefore may cause widespread alterations to synaptic signalling resulting in impairment of the function of the common core network which, in turn may explain the diverse acute and long-term behavioural and cognitive consequences of cannabis use(21,85,86).

Our second major finding was that the effect of THC on the pooled effect-size of regional brain signal was related to a proxy measure of regional CB1R density. The multiple linear regression model identified no significant relationship between CNR2 gene expression (a proxy measure of CB2R(36) with the effect size estimate). This is perhaps unsurprising as studies have shown that CB2 receptors are predominately distributed peripherally(87) with limited central distribution. Moreover, THC has less efficacy in its partial agonistic affinity to CB2 receptors compared with CB1 receptors in vitro(46). The brain regions found to be modulated by THC in our core analysis, including the anterior cingulate, amygdala, striatum, and cerebellum are known to be rich in CB1R(35). We show, for the first time, that a linear relationship exists between the effect of THC on increases in brain signal (as indexed by the pooled effect-size estimate) and CNR1 gene expression levels (as estimated on the basis of an average from 6 post-mortem brains of healthy individuals obtained from Allen Human Brain Atlas), a proxy measure of CB1R availability, across the whole brain(41). These findings are important as the CB1R is the main molecular target of THC in the human brain, where it has partial-agonist effects(46,88). Our findings thus provide novel —albeit indirect— evidence that the effects of THC on human brain function are in part related to local CB1 receptor availability, and complement independent experimental evidence that the acute effects of THC on human behaviour may be mediated by its effects on CB1R. See Supplementary Discussion 2 for additional discussion on CB1R mediating the effects of THC.

Our third key result was the identification of a relationship between THC dose and the effect-size estimates of activation signal across a range of brain substrates. We found a positive relationship between THC dose and its effects in the anterior cingulate cluster (comprising the dorsal and ventral regions), and a negative relationship in the supplementary motor area. These findings are significant as the anterior cingulate is believed have a role in social evaluation(89) and cognition(90), with functional alterations in individuals with high trait anxiety(91) and psychosis(92,93). Therefore, the dose-dependent effect of THC on the ventral cingulate may explain the findings of THC challenge studies(13,94) that investigated cognitive and psychological outcomes and have reported an association between higher doses of THC and increased psychotomimetic, anxiolytic, and cognitive impairments. Cannabis use has also been associated with motor impairments(95) with epidemiological reports suggesting a dose-related risk of motor vehicle accidents(96). However, one study has reported increased supplementary motor cortex activation with reduced psychomotor performance in chronic cannabis users during visual motor tasks(97). Interestingly, greater undirected functional connectivity between the dorsal anterior cingulate and supplementary motor area has been observed during proactive vs reactive motor control task conditions(98). Together, these findings suggest that the dose-response effects of THC on psychomotor dysfunction may, in part, be mediated by its effects on these brain regions, which could have implications for understanding how THC impairs the operation of heavy machinery in everyday life in cannabis users or patients prescribed THC-based medications. Emotional and cognition-agnostic effects of THC and its relationship with frontal cortical executive functioning as well as top-down control of subcortical structures are further discussed in Supplementary Discussion 3. Although, in our dose-response analyses, we identified the study by Battistella et al.,(74) as being a potential outlier, we refrained from excluding the study from dose-response association analyses in accordance with current thinking in this regard (please see further elaboration of this in Supplementary Discussion 4) and instead advise appropriate caution in the interpretation of the dose-response results.

### Limitations

The results presented here are to be considered in light certain key limitations. Firstly, our results are based on summary data from individual studies rather than individual participant level imaging data from the same participants carrying out multiple different cognitive and emotional processing tasks as well as actual baseline CB1R data in the same participants measured using PET imaging. This would have allowed more direct testing of our hypotheses. While future endeavours should aim to carry out such studies, conducting them in over 300 participants as reported herein is likely to be challenging both in terms of resources as well as logistics. The present meta-analysis, in contrast, provides an early insight into these questions using existing data. Another key caveat to be considered while interpreting our meta-analytic results is related to the issue of heterogeneity across the included studies. While this is inherent to any meta-analytic endeavour, our steps to examine the extent to which they may have influenced our results indicate that they are unlikely to have substantially affected our key conclusions. Limitations are discussed in greater detail in Supplementary Discussion, Methodological considerations & heterogeneity.

Notwithstanding these limitations, the three major findings of the current study extend previous evidence on the effects of THC to specifically link (a) the molecular effects of THC at the CB1 receptor to (b) its physiological (haemodynamic) effects on regional brain signal activation, which together may underlie (c) the acute cognitive and behavioural consequences of cannabis use. Only through meta-analytic synthesis of 22 studies across 372 participants in computational unison were we able to demonstrate that the pleiotropic effects of THC at each of these levels of observation may be related to its molecular target— the CB1 receptor. Here we present a potential mechanistic explanation for the pleiotropic effects of THC by reporting its effects on a ‘integrative core’ of brain regions engaged across diverse cognitive and emotional processes(37), where its effects are in turn related to the availability of its main central molecular target across the brain.

## Acknowledgments and Disclosures

SB is supported by grants from the Medical Research Council, UK; NIHR, UK; Wellcome trust; European Commission; Dowager Countess Eleanor Peel trust; Psychiatry Research trust; Parkinson’s UK; Rosetrees Trust; Alzheimer’s Research UK. SB has participated in advisory boards for or received honoraria as a speaker from Reckitt Benckiser, EmpowerPharm/SanteCannabis and Britannia Pharmaceuticals. All of these honoraria were received as contributions toward research support through King’s College London, and not personally. SB also has an ongoing collaboration with Beckley Canopy Therapeutics/ Canopy Growth (investigator initiated research) wherein they are supplying study drug for free for charity (Parkinson’s UK) and NIHR (BRC) funded research. MV is funded by the National Institute for Health Research Biomedical Research Centre at South London and Maudsley National Health Service Foundation Trust and King’s College London, and by the Wellcome Trust Digital Award 215747/Z/19/Z. MV has received consulting honoraria from GSK. The views expressed are those of the authors and not necessarily those of the NHS, the NIHR or the Department of Health.

