## Supplement for "Task-independent acute effects of delta-9-tetrahydrocannabinol on human brain function and its relationship with cannabinoid receptor gene expression: a neuroimaging meta-regression analysis"

***Supplement 1***

### *Supplementary Methods*

#### *Search Strategy*

Two categories of search terms were employed: 1) cannabis, marijuana, marihuana, THC, tetrahydrocannabinol and 2) fMRI, functional magnetic resonance, imaging, magnetic resonance, MRI, single photon emission tomography, SPECT, positron emission tomography, PET, spectroscopy, MRS, perfusion, blood flow, ASL. Categories were combined using Boolean Operator commands. Search of key terms was restricted to titles and abstracts within human studies.

#### *Eligibility Criteria*

Articles that did not assess the effects of THC-rich cannabis extract or THC on brain activity, or investigated connectivity were excluded, along with studies that primarily assessed psychological effects without any neuroimaging component. Studies that measured brain chemistry (such as PET studies investigating dopamine release or Magnetic Resonance Spectroscopy to measure glutamate) were not included.

We combined several imaging modalities as well as those with and without activation paradigms as we were interested in the distribution of effects across the brain rather than the specific effects associated with different cognitive activation tasks (e.g., n-back for working memory).

Two researchers independently performed the data search and extraction, with the final literature search conducted on 15/02/21. For any discrepancies that arose between the two researchers, a senior investigator was consulted.

#### ***Data Extraction***

Using the combined maps, the arithmetic mean of the effect size of the different functional tasks was calculated, while adjusting for variance by accounting for the number of tasks and the correlation among them, so that the combined study has the same statistical significance (z-value) as when using the effect size of the mean response. In other words, the effect size reflects the response of an individual study, and the fact that the combined study includes several tasks reduces the variance in a similar way that large samples do(1). Where maps were unavailable, whole-brain coordinates with their t-statistic were manually extracted from the published article for the conditions of interest (THC<PLB and THC>PLB). Where no t-statistic was reported, the given z-value or p-value was transformed into the t-value using a converter(2). Extracted data was cross-checked for errors between the two researchers.

For two studies(3,4), we had thresholded statistical maps, i.e., the voxels that originally had t-values below the threshold had been set to zero causing SDM to erroneously understand that the thresholded voxels are null. As in previous studies(5,6), to overcome this problem we used the following strategy: a) we created a grid mask image, that grouped the voxels in cubes; b) then multiplied the thresholded statistical maps by this mask, thus partitioning the unthresholded clusters in cubes; and c) finally identified the peak of each of these cubes. With this strategy, we obtained many local peaks (about 200 per image), and thus our analytic procedure (please see below) could recreate the unthresholded clusters with negligible error, while correctly imputing the effect sizes of the thresholded voxels.

#### ***Data Analysis***

We chose not to use SDM-PSI in for our analysis as we had already commenced the study when SDM-PSI was still very novel. $Q_H$ statistics were assessed in terms of a chi-square distribution and reported after conversion to standard z values to create a map. This map was overlaid on the final mean map for visual inspection of areas of overlapping significant heterogeneity with areas of thresholded activation or attenuation signal. For each significant cluster for THC vs placebo comparison, Egger's test was used to assess the asymmetry of funnel plots to examine potential publication bias(7). This was conducted to assess whether small or imprecise studies reported larger effect sizes. 'Small' studies with negative results may be less likely to be published, and conversely, large studies are more likely to be published even with negative findings. Coordinates of cluster peaks and cluster extent are reported using MNI coordinates. Each cluster peak was examined using a human brain atlas in order to visually inspect the peak region(8). In a jack-knife leave-one-out sensitivity analysis, each analysis was repeated excluding 1 single study at a time to establish whether each cluster remained statistically significant.

***Whole brain correlation with CNR1 and CNR2 gene expression***

The Allen Human Brain Atlas contains microarray expression data in 3,702 spatially distinct tissue samples collected from six neurotypical adult human brains(9). The samples are distributed across cortical, subcortical, brainstem and cerebellar regions, quantifying the expression of more than 20,000 genes.

First, transcriptional gene expression data from microarray probes were downloaded from the Allen Human Brain Atlas database. Microarray probe-to-gene reannotation was conducted with information from Arnatkevičiūtė et al., 2019(10). Probes that could not be reliably matched to genes were discarded. We then performed intensity-based filtering of probes to remove those that did not exceed background noise using a threshold of 0.5 (i.e., probes must exceed background noise in 50% of all tissue samples). Following this, representative probes, with the highest differential stability (i.e., the probe with the most consistent spatial pattern across the donor brains), in indexing the cannabinoid-receptor 1 (CNR1) and cannabinoid-receptor 2 (CNR2) genes coding for the CB1R and CB2Rs respectively, were selected. Microarray samples were then matched to 78 cortical regions, parcellated across the Desikan–Killiany brain atlas(11). For precise sample to region matching, we initially determined if the microarray sample fell directly within a region of the atlas. If the sample did not, they were assigned to a region within a distance of 2mm of the Desikan-Killany parcels. In cases where there were multiple nearby regions, the closest region was determined by calculating the centre-of-mass of the abutting regions. We then transformed expression values for the CNR1 and CNR2 genes across samples, for each donor, using a scaled robust sigmoid normalisation function. Finally, expression values for microarray samples, assigned to each region of the Desikan-Killany atlas, were averaged independently for each donor and then averaged across donors.

### ***Supplementary Results:***

#### ***Funnel Plots For The Blobs In Core Analysis With Egger's Test For Asymmetry And Jack-Knife Leave One Out Analysis Matrix***

The funnel plots below depict Hedge's g effect size estimates from individual studies (x axis) against the standard error (SE) (y axis). Larger, more powerful studies are placed at the top of the funnel plot. The effect size estimates from smaller studies are distributed at the bottom of the plot, with the spread narrowing among larger studies(12). In the absence of bias and between study heterogeneity, the scatter should arise from variations in sampling shown by a symmetrical inverted funnel(13). Present within the triangle should be 95% of the studies if there is no bias and the fixed effect assumption is true (the true effect of THC is the same in each study). Factors that may cause asymmetry in the funnel plot may be heterogeneity, reporting bias, choice of statistics being plotted, and chance(13). Egger's test is a statistical test that was used to quantitatively asses the asymmetry of funnel plots(7). The Egger's test suggested potential publication bias for 3 clusters: the right superior frontal gyrus (THC>PLB; Egger's test P= 0.044; peak coordinates 6, 62, -4), the right insula (THC<PLB; Egger's test P=0.037; peak coordinates -44, -12, 8), and the right Rolandic operculum (THC<PLB; Egger's test P= 0.044; peak coordinates 48, -8, 10).

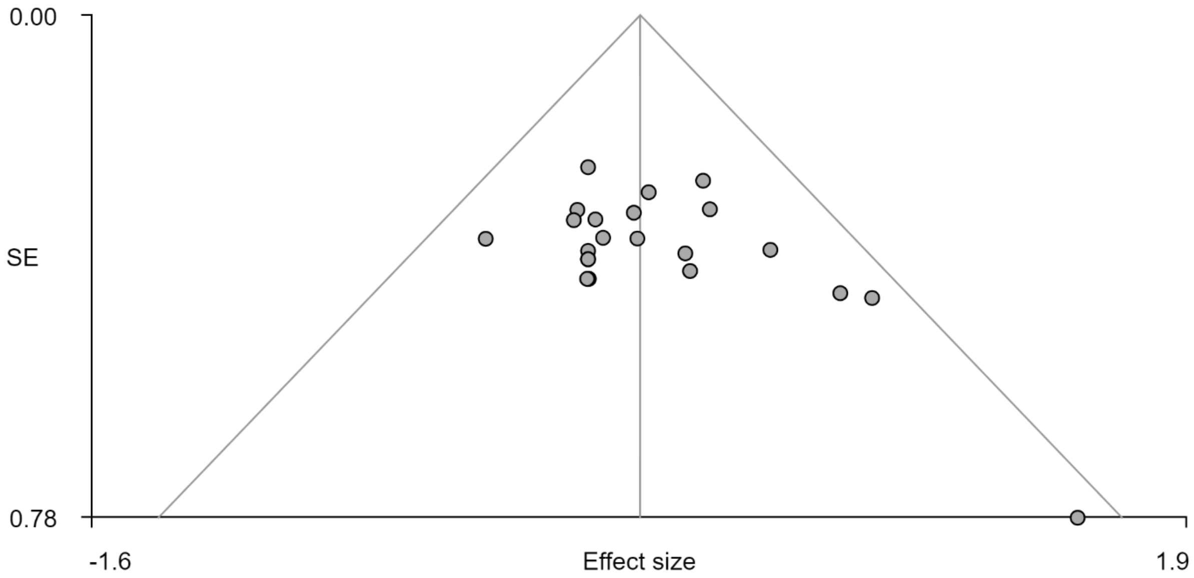


*Supplementary Figure 1. THC>placebo, right superior frontal gyrus, Egger's test, bias= 1.73, t: 2.14, df: 21, p: 0.044*

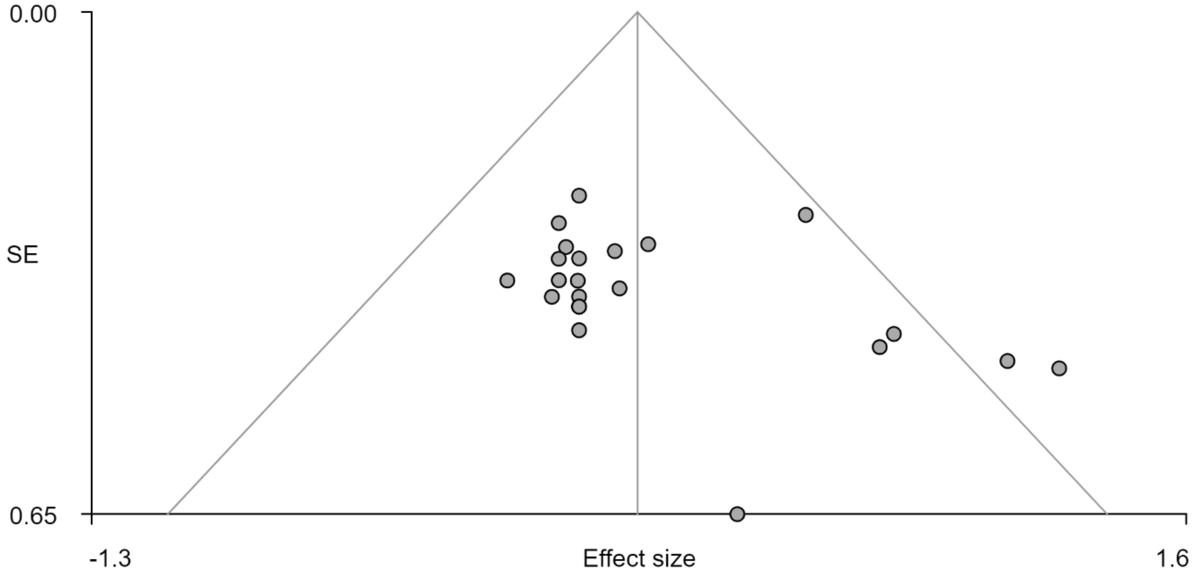


*Supplementary Figure 2. THC>placebo right superior frontal gyrus, Egger's test, bias= 2.03, t: 1.93, df: 21, p: 0.067*

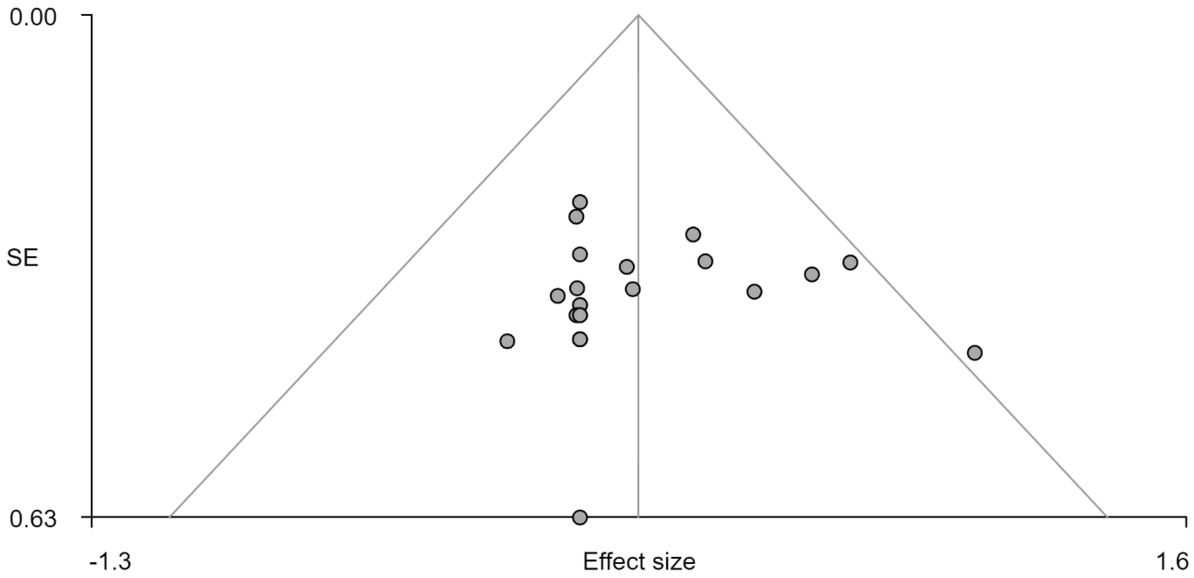


*Supplementary Figure 3. THC>placebo, right middle temporal gyrus Egger's test, bias= -0.05, t: -0.05, df: 21, p: 0.961*

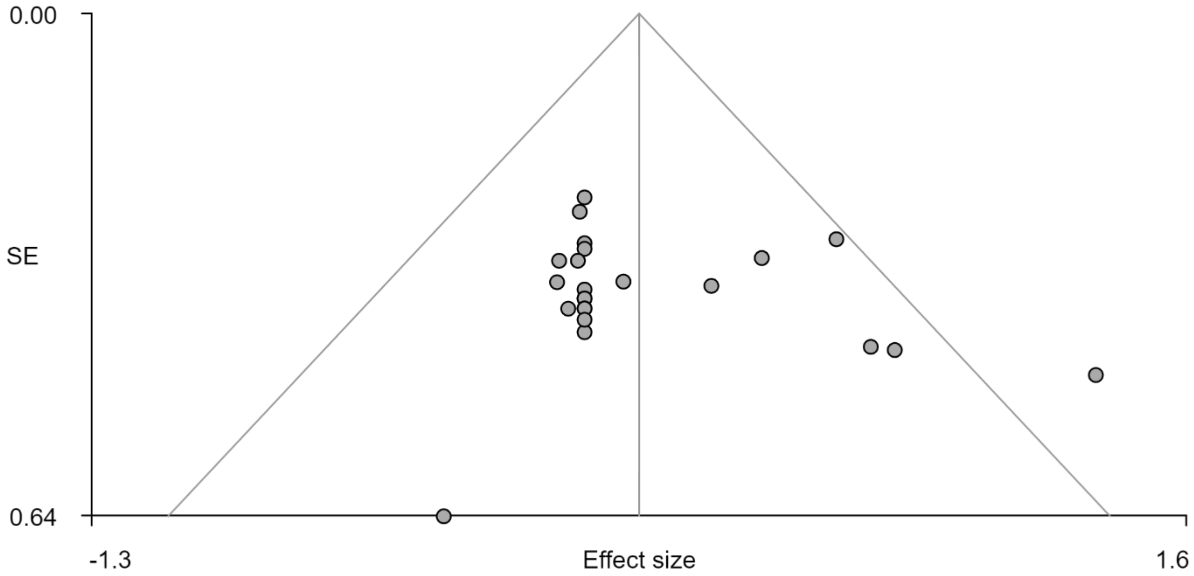


*Supplementary Figure 4. THC>placebo, R cerebellum crus II, Egger's test, bias= 1.10, t: 1.05, df: 21, p: 0.303*

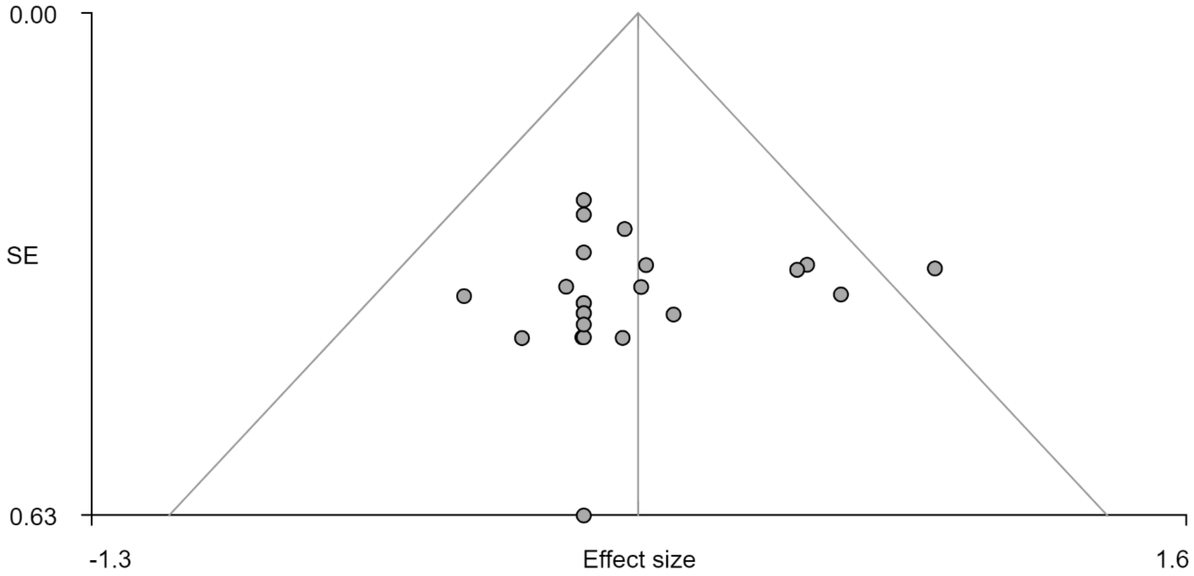


*Supplementary Figure 5. THC>placebo, right inferior occipital gyrus, Egger's test, bias= -0.36, t: -0.36, df: 21, p: 0.720*

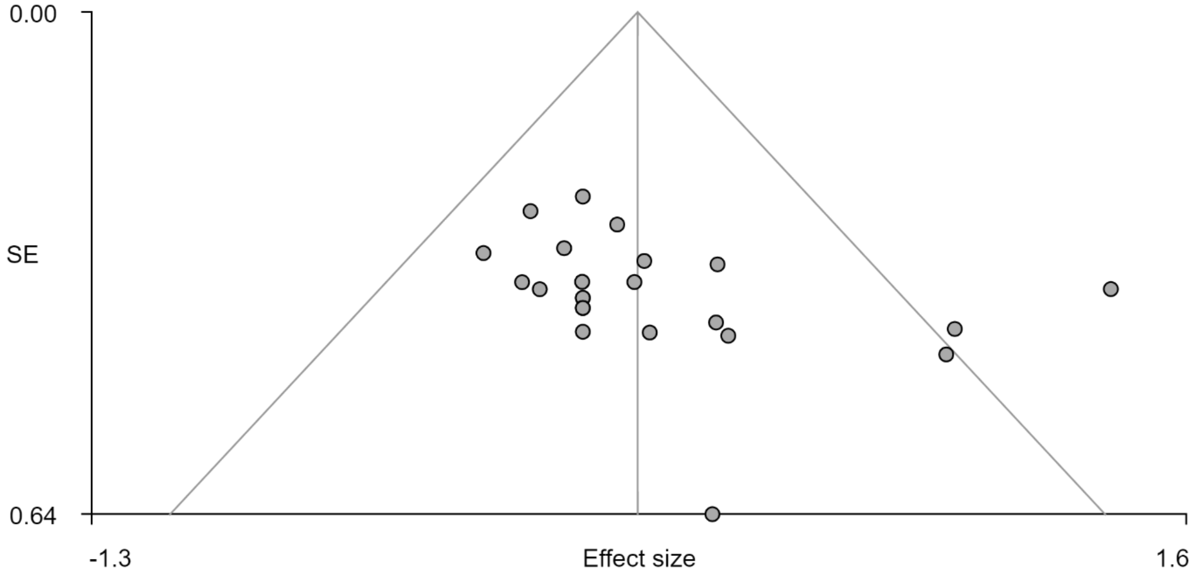


*Supplementary Figure 6. THC>placebo, right amygdala, Egger's test, bias= 2.24, t: 1.92, df: 21, p: 0.069*

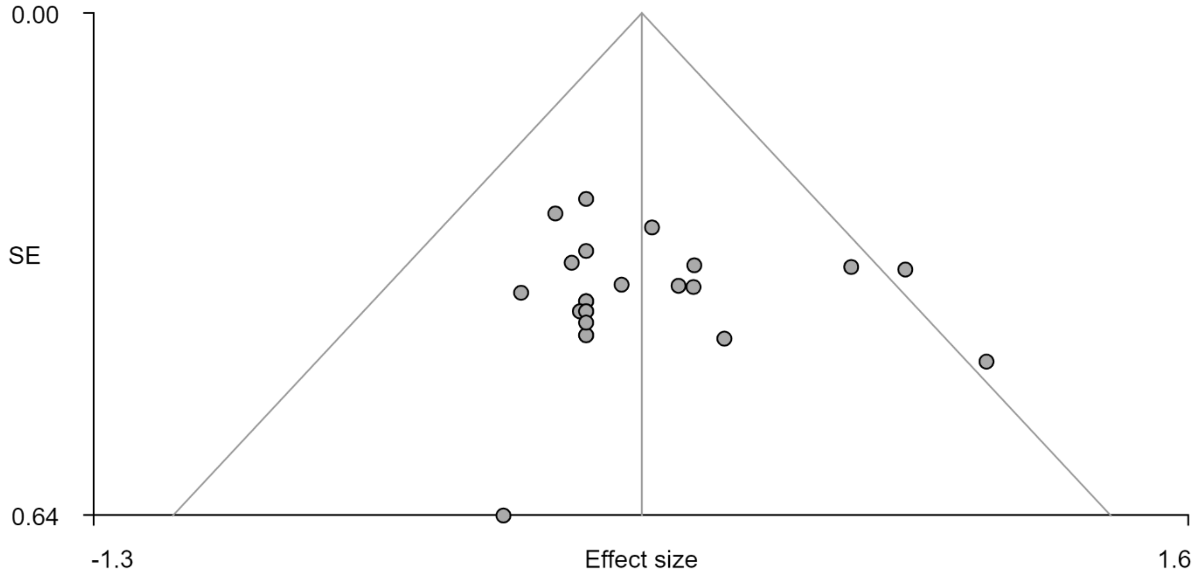


*Supplementary Figure 7. THC>placebo, left cerebellum lobule VIIB, Egger's test, bias= 0.45, t: 0.45, df: 21, p: 0.654*

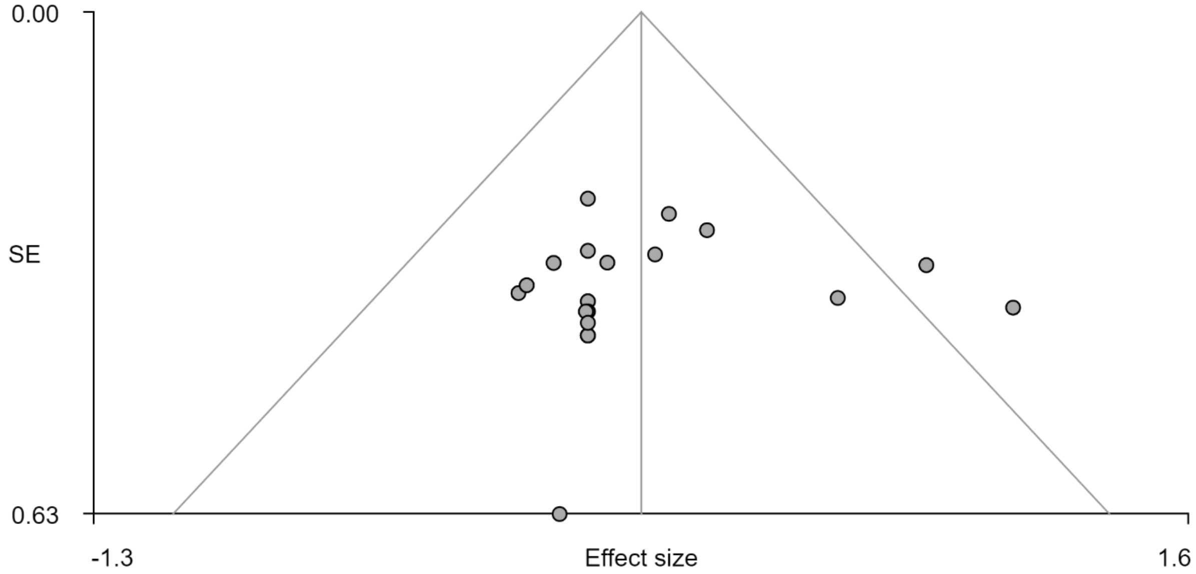


*Supplementary Figure 8. THC>placebo, left precentral gyrus, Egger's test, bias= -0.44, t: -0.41, df: 21, p: 0.688*

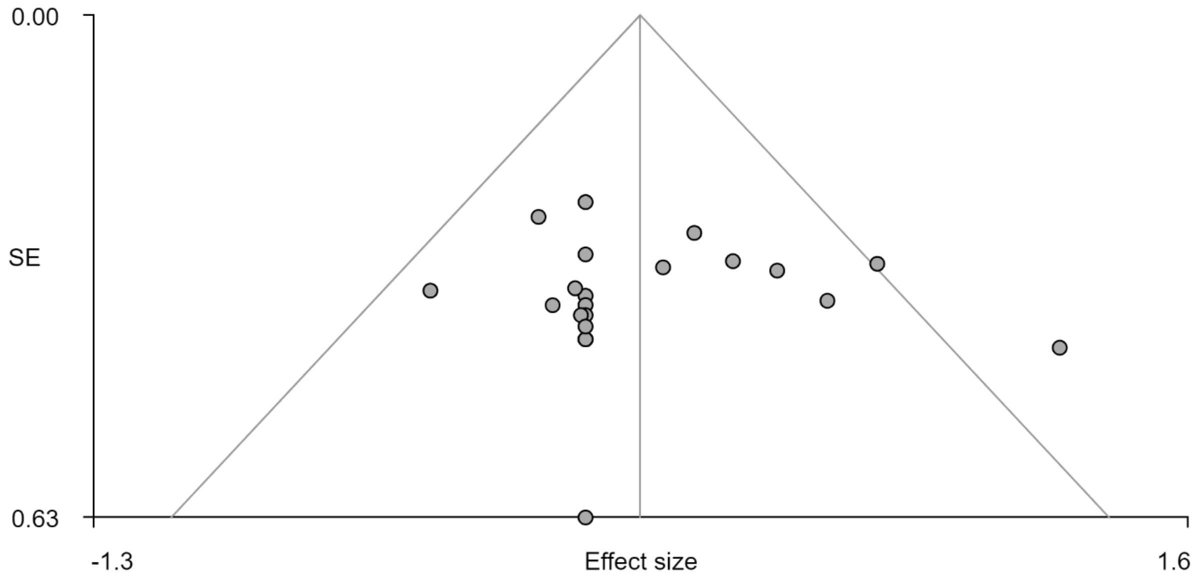


*Supplementary Figure 9. THC>placebo, left thalamus, Egger's test, bias= 0.27, t: 0.24, df: 21, p: 0.814*

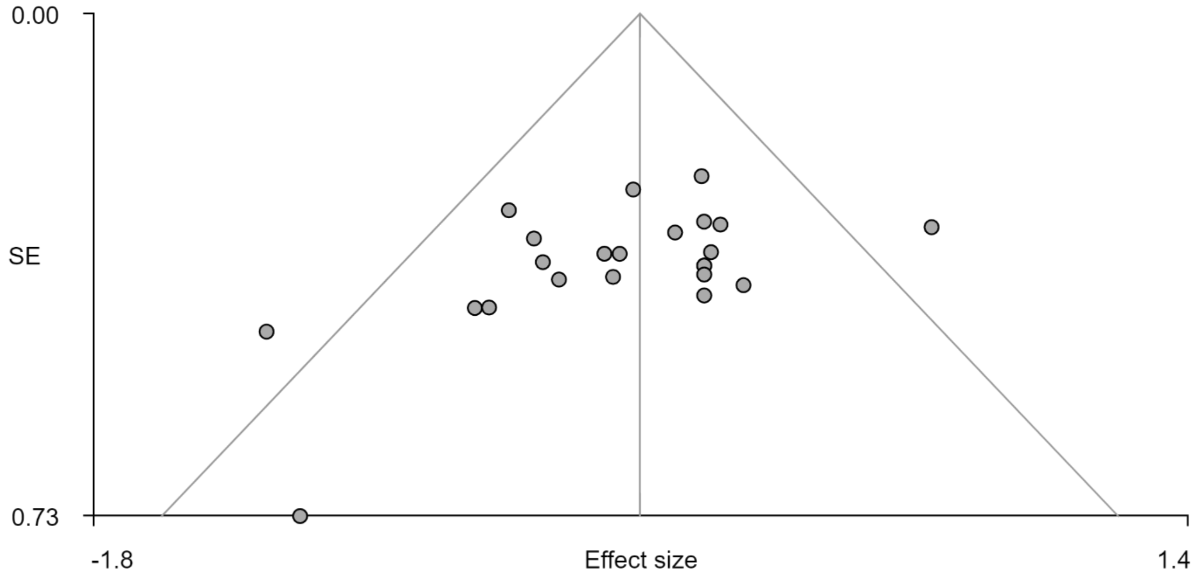


*Supplementary Figure 10. THC<placebo, left insula, Egger's test, bias= -2.33, t: -2.23, df: 21, p: 0.037*

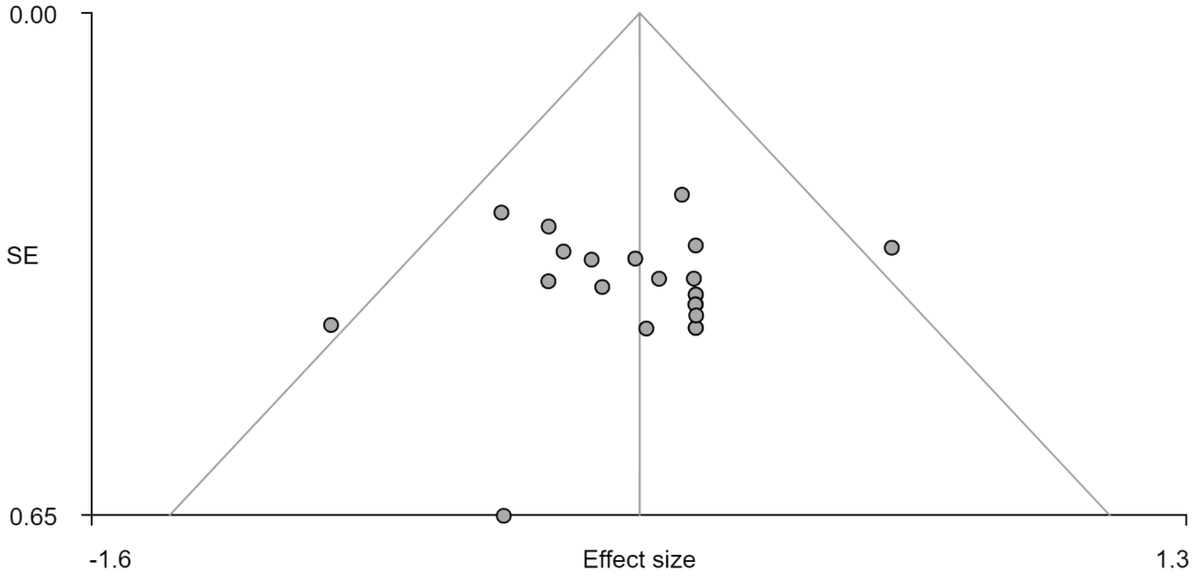


*Supplementary Figure 13. THC<placebo, left precentral gyrus, Egger's test, bias= -0.04, t: -0.04, df: 21, p: 0.971*

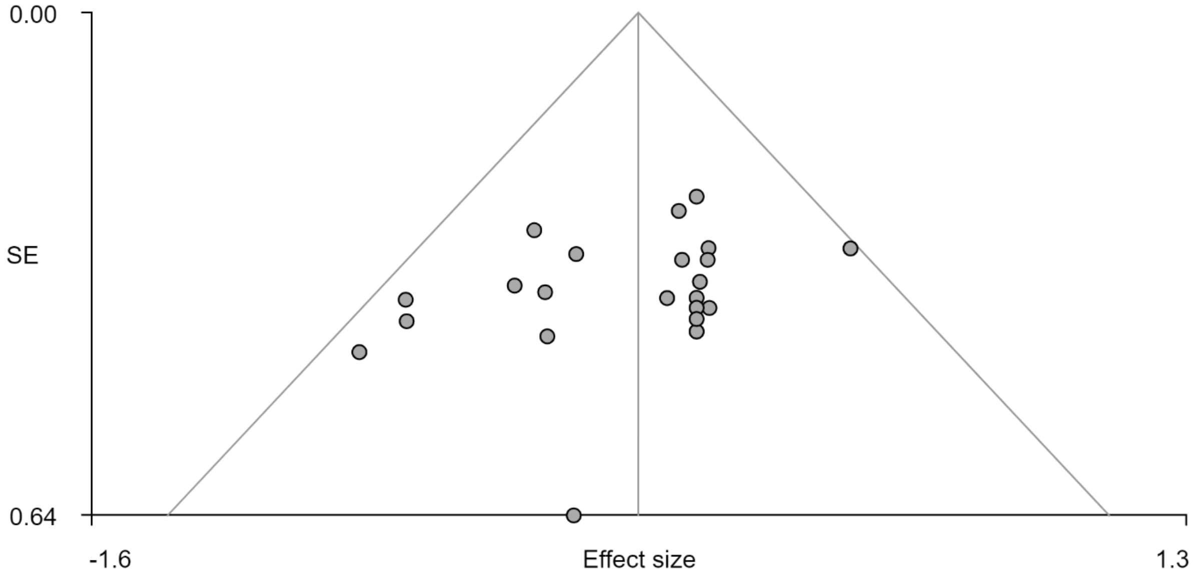


*Supplementary Figure 14. THC<placebo, right superior temporal gyrus, Egger's test, bias= -1.48, t: -1.56, df: 21, p: 0.135*

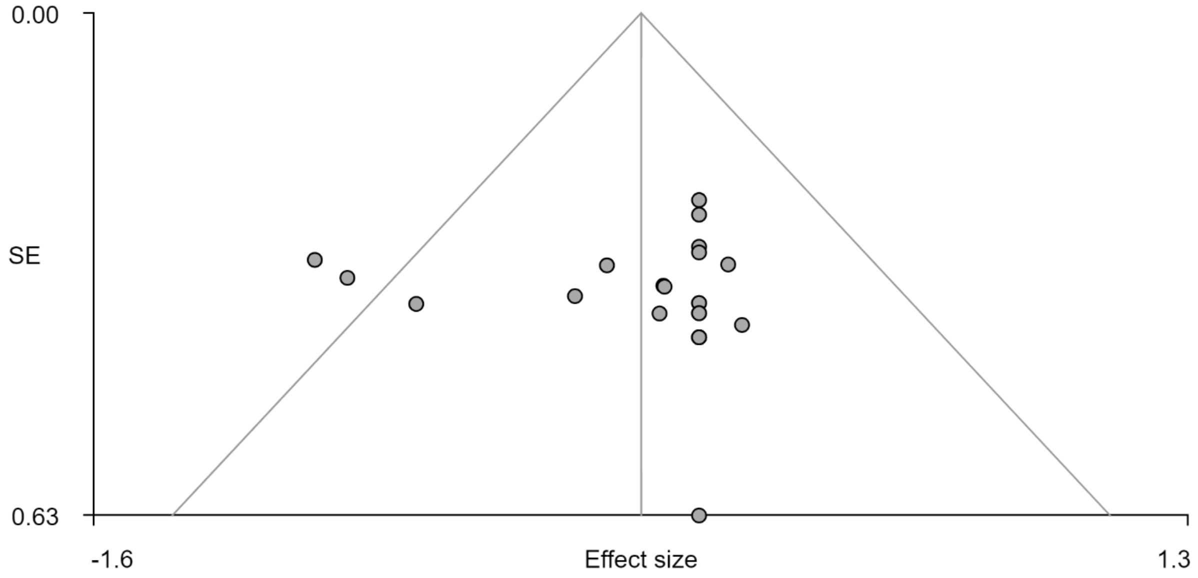


*Supplementary Figure 15. THC<placebo, left angular gyrus, Egger's test, bias= 0.14, t: 0.12, df: 21, p: 0.902*

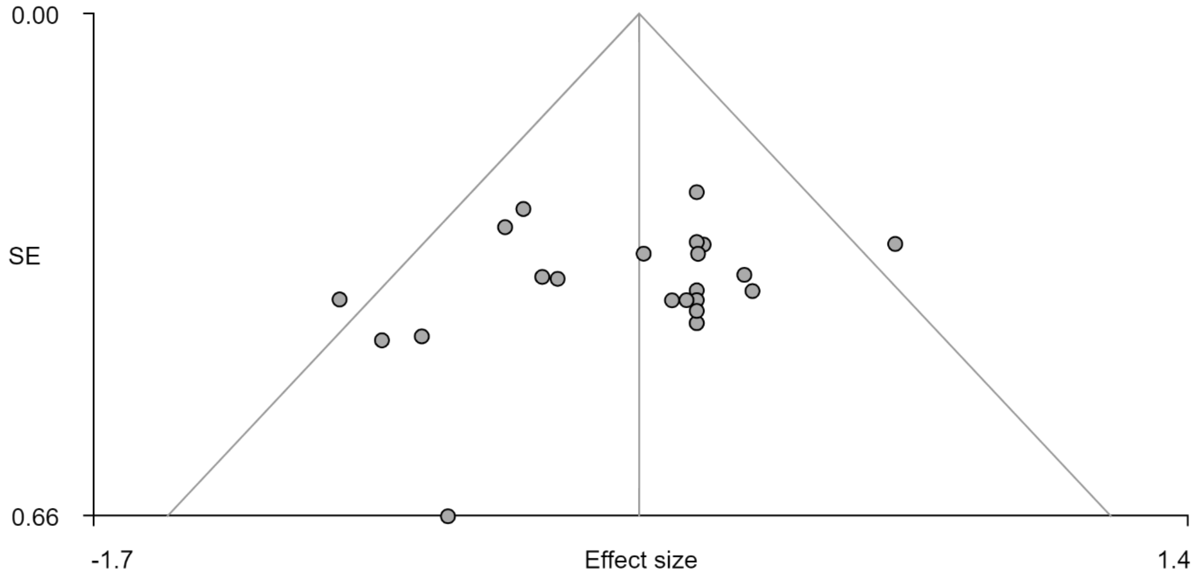


*Supplementary Figure 16. THC<placebo, cerebellum lobule IV/V, Egger's test, bias= -1.15, t: -1.02, df: 21, p: 0.318*

*Supplementary Table 1. Jack-knife leave-one-out sensitivity analysis, each analysis was repeated excluding 1 single study at a time to establish whether each cluster remained statistically significant. 87% of all clusters survived this analysis indicated with dots. Crosses indicate regions where clusters did not survive.*

| | *R superior frontal gyrus* | *R superior frontal gyrus* | *R middle temporal gyrus* | *R cerebellum crus II* | *R inferior occipital gyrus* | *R amygdala* | *L cerebellum lobule VIIB* | *L precentral gyrus* | *L thalamus* | *L insula* | *R rolandic operculum* | *R cuneus cortex* | *L precentral gyrus* | *R superior temporal gyrus* | *L angular gyrus* | *Cerebellum lobule IV/V* |
|---|---|---|---|---|---|---|---|---|---|---|---|---|---|---|---|---|
| *Battistella*(3) | ● | ● | ● | ● | ● | ● | ● | × | ● | ● | ● | ● | × | ● | ● | × |
| *Bhattacharyya*(14) | ● | ● | ● | ● | ● | ● | ● | ● | ● | ● | ● | ● | ● | ● | ● | ● |
| *Bhattacharyya*(15) | ● | ● | ● | ● | ● | ● | ● | ● | ● | ● | ● | ● | ● | ● | ● | ● |
| *Bhattacharyya*(16) | ● | ● | ● | ● | ● | × | ● | ● | × | ● | ● | ● | ● | ● | ● | ● |
| *Bhattacharyya*(17) | ● | ● | ● | ● | ● | ● | ● | ● | ● | ● | ● | ● | ● | ● | ● | ● |
| *Bossong*(18) | ● | ● | ● | × | ● | ● | ● | × | × | ● | ● | ● | ● | ● | × | ● |
| *Bossong*(19) | ● | ● | ● | ● | ● | ● | × | ● | × | ● | ● | ● | ● | ● | × | ● |
| *Bossong*(20) | ● | ● | ● | ● | ● | ● | ● | × | ● | ● | ● | ● | ● | ● | ● | ● |
| *Bossong*(21) | ● | ● | ● | ● | ● | × | ● | ● | ● | ● | ● | ● | ● | ● | ● | ● |
| *Bossong*(22) | ● | ● | ● | ● | ● | ● | ● | ● | × | ● | ● | ● | ● | ● | ● | ● |
| *Freeman*(4) | ● | ● | ● | ● | ● | ● | ● | ● | ● | ● | ● | ● | ● | ● | ● | × |
| *Jansma*(23) | ● | ● | ● | ● | ● | ● | ● | × | × | ● | ● | ● | ● | × | × | ● |
| *Lee*(24) | ● | ● | ● | ● | ● | ● | ● | × | ● | ● | ● | ● | ● | ● | × | ● |
| *O'Leary*(25) | ● | ● | ● | ● | ● | × | ● | × | ● | ● | ● | ● | ● | ● | × | ● |
| *O’Leary*(26) | ● | ● | ● | ● | ● | × | ● | × | ● | ● | ● | ● | ● | ● | × | × |
| *O'Leary*(27) | ● | ● | ● | ● | ● | ● | × | × | ● | ● | ● | ● | ● | × | × | × |
| *O'Leary*(28) | ● | ● | ● | ● | ● | ● | ● | × | ● | ● | ● | ● | ● | ● | × | ● |
| *Rabinak*(29) | ● | ● | ● | ● | ● | ● | ● | × | ● | ● | ● | ● | × | ● | × | ● |
| *van Hell*(30) | ● | ● | ● | ● | ● | ● | ● | × | ● | ● | ● | ● | ● | ● | × | ● |
| *Walter*(31) | ● | ● | ● | ● | ● | ● | ● | ● | ● | ● | ● | ● | ● | ● | ● | ● |
| *Walter*(31) | ● | ● | ● | ● | ● | ● | ● | ● | ● | ● | ● | ● | ● | ● | ● | ● |
| *Walter*(32) | ● | ● | ● | ● | ● | × | ● | × | ● | ● | ● | ● | ● | ● | × | ● |
| *Winton-Brown*(33) | ● | ● | ● | ● | ● | ● | ● | ● | ● | ● | ● | ● | ● | × | ● | ● |

***Supplementary subgroup analyses***

We performed subgroup analysis on studies that employed similar cognitive paradigms to investigate the differential effect of THC on brain activation.

### 3.4 Subgroup analyses

*Emotion processing*

Within the emotional processing subgroup, 42 participants were compared in a crossover design following THC compared with placebo while processing emotional pictures(16,19,29).

*Supplementary Table 2. Areas of increased and attenuated brain signal following THC, compared with placebo, during emotional processing stimuli*

| *Comparison* | *MNI coordinate* | | | *SDM-Z* | *P* | *Voxels* | *Region* |
|---|---|---|---|---|---|---|---|
| | x | y | z | | | | |
| *THC>PLB* | 34 | -18 | -4 | 2.105 | <0.001 | 341 | R putamen (extending to: R insula, R striatum, R heschl gyrus) |
| | 12 | -24 | -2 | 2.124 | <0.001 | 187 | R thalamus (extending to: R hippocampus, R thalamus) |
| | 20 | 20 | -16 | 1.891 | <0.001 | 85 | R superior frontal gyrus, orbital part (extending to: R striatum, R gyrus rectus, R olfactory cortex) |
| | 56 | -26 | 16 | 1.628 | 0.003 | 21 | R superior temporal gyrus (extending to: R supramarginal gyrus) |
| | 28 | -36 | 10 | 2.071 | <0.001 | 17 | Caudate (extending to: R hippocampus) |
| | 44 | -34 | 4 | 1.527 | 0.003 | 14 | Superior temporal gyrus |
| | 48 | 8 | -12 | 1.661 | 0.002 | 13 | R temporal pole, superior temporal gyrus |
| *THC<PLB* | -44 | -68 | 38 | -2.169 | <0.001 | 330 | L angular gyrus (extending to: L middle temporal gyrus) |
| | -40 | -28 | 46 | -2.129 | <0.001 | 241 | L postcentral gyrus (extending to: inferior parietal [excluding supramarginal and angular] gyri) |
| | -4 | 24 | 58 | -2.157 | <0.001 | 221 | L supplementary motor area (extending to: L medial superior frontal gyrus) |
| | -60 | 8 | 12 | -1.985 | <0.001 | 101 | L inferior frontal gyrus, opercular part (extending to: pre and postcentral gyri) |
| | -8 | -54 | 72 | -1.933 | 0.001 | 16 | L precuneus (extending to: L superior parietal gyrus) |
| | -14 | -42 | 78 | -2.046 | <0.001 | 16 | L postcentral gyrus (extending to: L precuneus and L paracentral gyri) |
| | -8 | -40 | -8 | -1.774 | 0.003 | 15 | L cerebellum, hemispheric lobule IV / V |

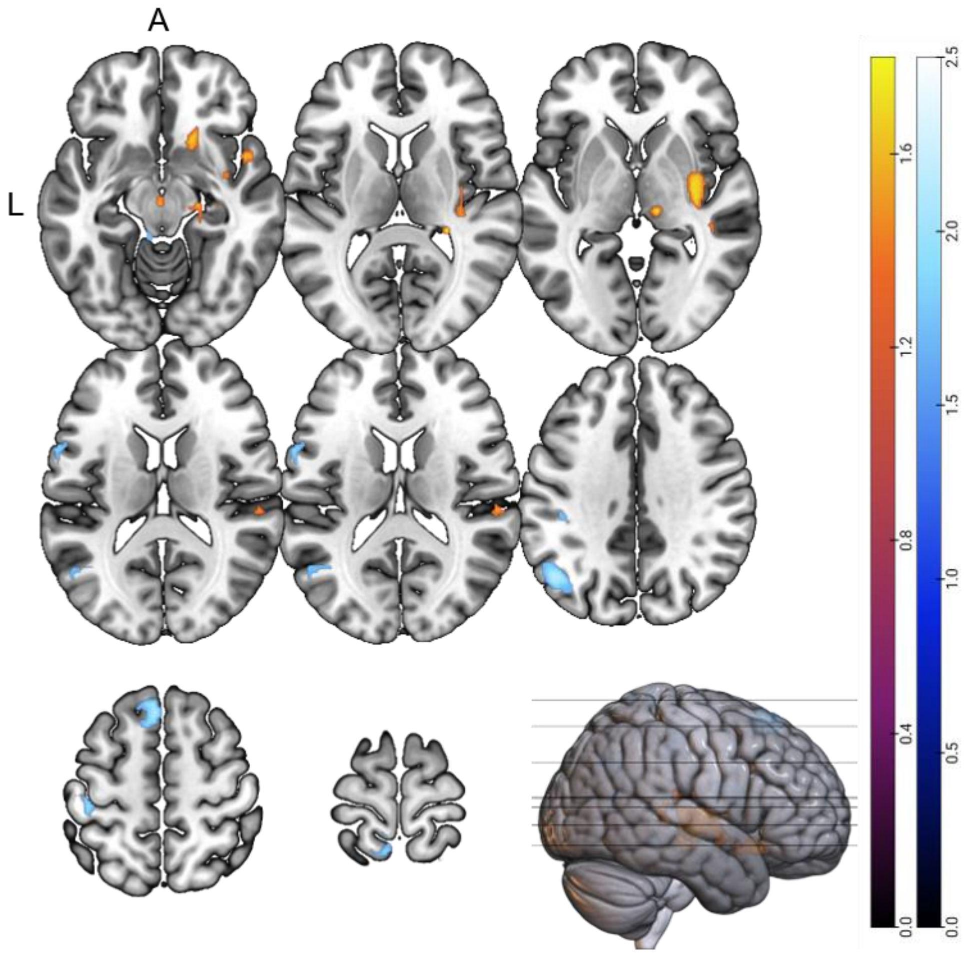


*Supplementary Figure 17. Emotional processing: a) Areas of increased activation signal after THC compared with placebo, b) Areas of attenuated activation signal after THC compared with placebo*

#### *Reward paradigms*

Within the reward processing subgroup, 37 participants were compared in a crossover design following THC compared with placebo while performing the MIDT(23,30) and listening to musically rewarding stimuli(4).

*Supplementary Table 3. Areas of increased and attenuated activation signal following THC, compared with placebo, during reward processing stimuli*

| *Comparison* | *MNI coordinate* | | | *SDM-Z* | *P* | *Voxels* | *Region* |
|---|---|---|---|---|---|---|---|
| | x | y | z | | | | |
| *THC>PLB* | -24 | -4 | 44 | 1.829 | <0.001 | 155 | L middle frontal gyrus (extending to: L precentral gyrus, L superior dorsolateral frontal gyrus) |
| | 8 | 44 | 30 | 1.957 | <0.001 | 142 | R median cingulate / paracingulate gyri (extending to: anterior cingulate gyrus and medial superior frontal gyrus) |
| | 18 | 68 | 0 | 2.41 | <0.001 | 78 | R superior frontal gyrus, dorsolateral |
| | 12 | 58 | -8 | 2.344 | <0.001 | 57 | R superior frontal gyrus, medial orbital (extending to: R striatum) |
| | -40 | -64 | 8 | 1.733 | 0.001 | 41 | L middle temporal gyrus (extending to: L middle occipital gyrus) |
| | -16 | 12 | 70 | 1.692 | 0.002 | 42 | L superior frontal gyrus, dorsolateral (extending to: L middle frontal gyrus) |
| | -8 | -44 | -60 | 1.811 | 0.001 | 24 | Cerebellum |
| | -4 | -56 | -52 | 1.68 | 0.002 | 14 | L cerebellum, hemispheric lobule IX |
| *THC<PLB* | 40 | -32 | -16 | -4.284 | <0.001 | 176 | R parahippocampal gyrus/ fusiform gyrus |
| | -60 | -44 | 12 | -3.008 | <0.001 | 115 | L superior temporal gyrus (extending to: L middle temporal gyrus) |
| | 16 | -26 | 40 | -3.148 | <0.001 | 111 | R median cingulate / paracingulate gyri |
| | -54 | 4 | -4 | -3.037 | <0.001 | 109 | L superior temporal gyrus (extending to: L rolandic operculum, L temporal pole, L heschl gyrus) |
| | 48 | 12 | -28 | -3.207 | <0.001 | 83 | R temporal pole, middle temporal gyrus (extending to: R superior temporal gyrus) |
| | -8 | -104 | 0 | -3.096 | <0.001 | 64 | L middle occipital gyrus (extending to: L superior occipital gyrus) |
| | 32 | 8 | 54 | -2.847 | 0.001 | 56 | R middle frontal gyrus (extending to:R precentral gyrus) |
| | 52 | -28 | 48 | -2.999 | <0.001 | 36 | R postcentral gyrus (extending to: R inferior parietal [excluding supramarginal and angular] gyri) |
| | -48 | 16 | 44 | -2.589 | 0.002 | 33 | L middle frontal gyrus |
| | -12 | 44 | -10 | -2.527 | 0.002 | 16 | L superior frontal gyrus, medial orbital |
| | 64 | -28 | -8 | -2.498 | 0.003 | 16 | R middle temporal gyrus |
| | -56 | -28 | 20 | -2.419 | 0.004 | 10 | L supramarginal gyrus |

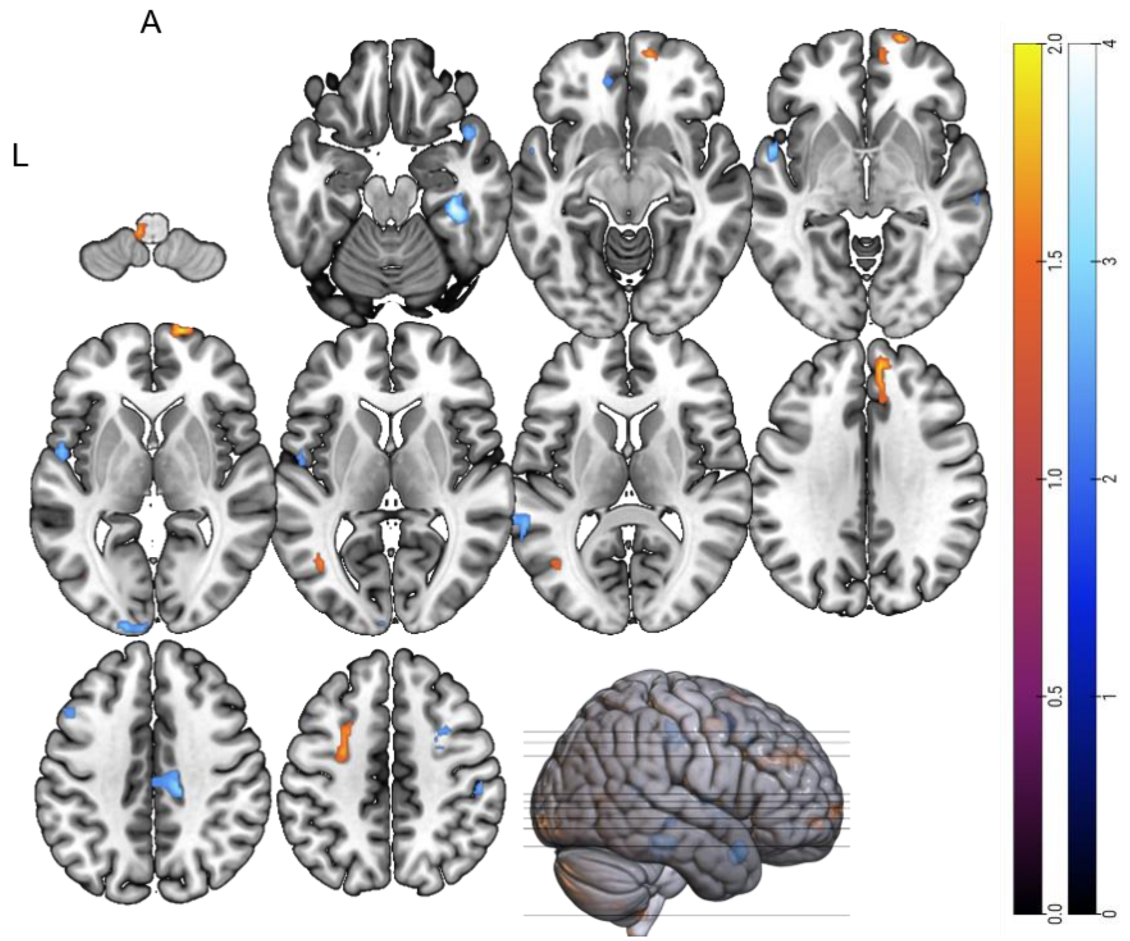


*Supplementary Figure 18. Reward processing: a) Areas of increased activation signal after THC compared with placebo, b) Areas of attenuated activation signal after THC compared with placebo*

*Sensory stimulation paradigms*

Within the sensory stimulation subgroup, 64 participants were compared in a crossover design following THC compared with placebo(24,31–33)

*Supplementary Table 4. Areas of increased and attenuated activation signal following THC, compared with placebo, during sensory processing stimuli*

| *Comparison* | *MNI coordinate* | | | *SDM-Z* | *P* | *Voxels* | *Region* |
|---|---|---|---|---|---|---|---|
| | x | y | z | | | | |
| *THC>PLB* | 4 | -24 | 28 | 1.67 | 0.004 | 73 | R mid cingulum (extending to R median cingulate/ paracingulate gyri) |
| | 52 | -42 | -6 | 2.05 | 0.001 | 37 | R middle temporal gyrus (extending to R inferior frontal gyrus) |
| | 14 | -86 | -10 | 1.69 | 0.004 | 25 | R lingual gyrus |
| | 26 | -82 | -4 | 1.70 | 0.003 | 14 | R fusiform gyrus (extending to R lingual gyrus) |
| | 52 | -42 | 44 | 1.67 | 0.004 | 14 | R supramarginal gyrus (extending to R inferior parietal gyri) |
| *THC<PLB* | -12 | 16 | 12 | -1.72 | <0.001 | 77 | L caudate nucleus |

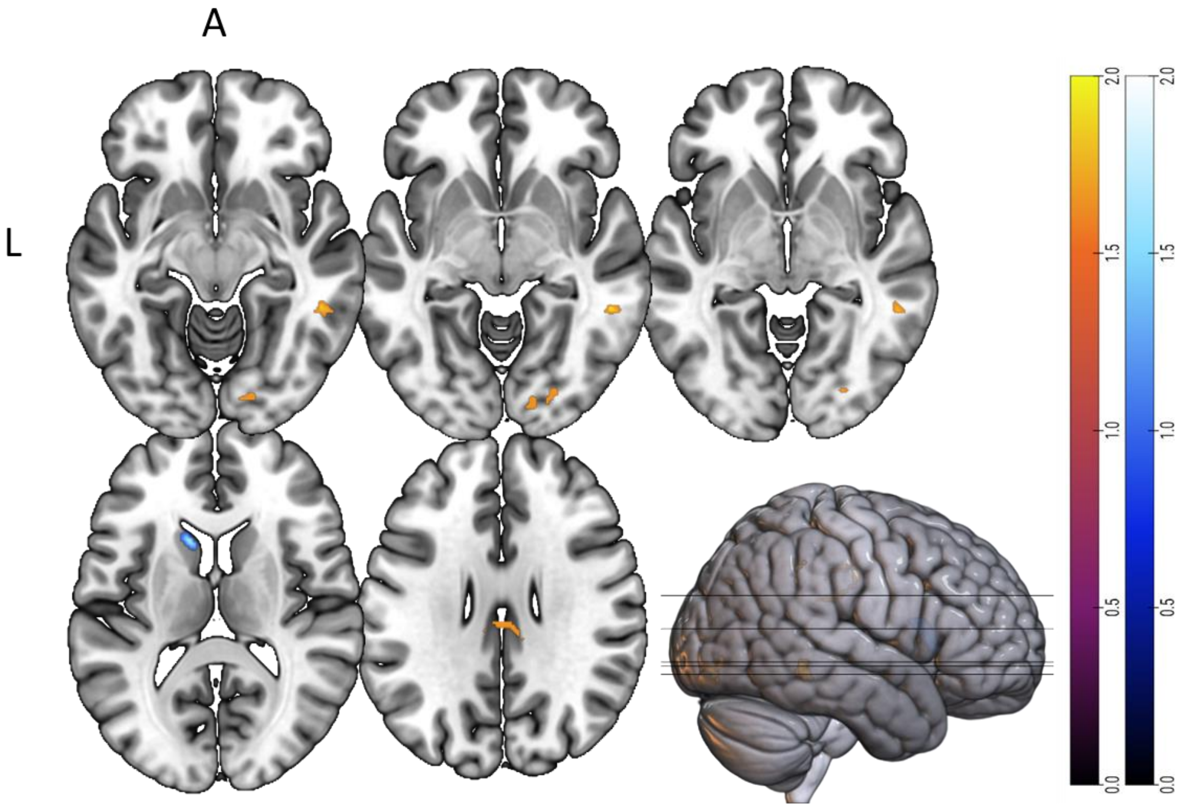


*Supplementary Figure 19. Sensory processing: a) Areas of increased activation signal after THC compared with placebo, b) Areas of attenuated activation signal after THC compared with placebo*

*Memory paradigms*

Within the memory processing subgroup, 41 participants were compared in a crossover design following THC compared with placebo while performing the verbal paired associates task(15), Sternberg item recognition(18), and the associative memory task(21).

*Supplementary Table 5. Areas of increased and attenuated activation signal following THC, compared with placebo, during memory processing*

| Comparison | MNI coordinate | | | SDM-Z | P | Voxels | Region |
|---|---|---|---|---|---|---|---|
| | x | y | z | | | | |
| *THC>PLB* | -28 | 48 | 22 | 2.169 | <0.001 | 253 | L middle frontal gyrus (extending to L superior frontal gyrus, dorsolateral & medial) |
| | -34 | -56 | -40 | 2.352 | <0.001 | 100 | L cerebellum |
| | 32 | 24 | 54 | 2.099 | 0.001 | 23 | R middle frontal gyrus (extending to R superior frontal gyrus, dorsolateral) |
| | -14 | 40 | 16 | 1.996 | 0.002 | 20 | L anterior cingulate cortex (extending to L medial superior frontal gyrus) |
| | -52 | 36 | 2 | 2.165 | <0.001 | 19 | L inferior frontal gyrus, triangular part |
| | 28 | 48 | 16 | 1.87 | 0.003 | 17 | R middle frontal gyrus (extending to R superior frontal gyrus, dorsolateral) |
| | 48 | -72 | 28 | 1.959 | 0.002 | 16 | R middle occipital gyrus (extending to R middle temporal gyrus) |
| | 44 | 16 | 32 | 1.875 | 0.002 | 13 | R inferior frontal gyrus, opercular part |
| | -12 | 44 | 40 | 1.897 | 0.002 | 13 | L superior frontal gyrus, dorsolateral |
| | -24 | -64 | -56 | 2.241 | <0.001 | 10 | L cerebellum, hemispheric lobule VIII |
| *THC<PLB* | -48 | -6 | -4 | -2.397 | 0.002 | 350 | L superior temporal gyrus (extending to L rolandic operculum, L heschl gyrus, L superior temporal gyrus, L insula, L supramarginal gyrus, L postcentral gyrus) |
| | 12 | -40 | 0 | -3.11 | <0.001 | 223 | R lingual gyrus (extending to R precuneus, L thalamus, R parahippocampal gyrus, R hippocampus, cerebellum vermic lobule IV/V) |
| | 4 | -48 | 56 | -2.397 | 0.002 | 149 | R precuneus (extending to L precuneus, R superior parietal gyrus, R paracentral lobule, R postcentral gyrus, L precuneus) |
| | 4 | -72 | 24 | -2.397 | 0.002 | 140 | R cuneus cortex (extending to R precuneus, R superior occipital gyrus, R superior parietal gyrus, L precuneus) |
| | -8 | -32 | -16 | -2.772 | <0.001 | 98 | L cerebellum (extending to L fusiform gyrus, L pons, L parahippocampal gyrus, L hippocampus, L cerebellum hemispheric lobule III) |
| | -30 | -28 | -6 | -2.391 | 0.002 | 43 | L hippocampus (extending to L parahippocampal gyrus) |
| | -14 | -58 | 58 | -2.388 | 0.002 | 42 | L precuneus |
| | 10 | -58 | -50 | -2.388 | 0.002 | 34 | R cerebellum, hemispheric lobule IX |
| | -52 | 6 | 16 | -2.366 | 0.002 | 16 | L precentral gyrus (extending to L inferior frontal gyrus, opercular part) |
| | -56 | -22 | 44 | -2.385 | 0.002 | 15 | L inferior parietal (excluding supramarginal and angular) gyri |
| | 22 | -68 | 54 | -2.388 | 0.002 | 14 | R superior parietal gyrus |

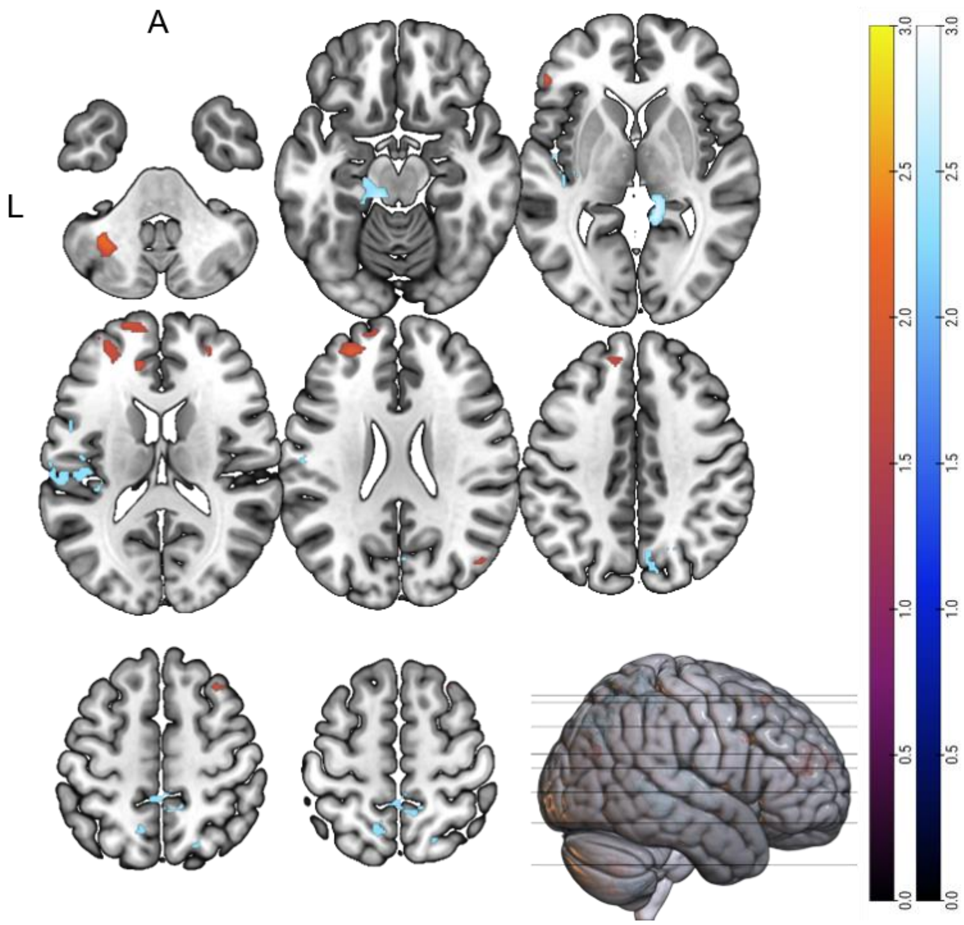


*Supplementary Figure 20. Memory processing: a) Areas of increased activation signal after THC compared with placebo, b) Areas of attenuated activation signal after THC compared with placebo*

#### *Attentional salience paradigms*

Within the attentional salience subgroup, 95 participants were compared in a crossover design following THC compared with placebo while performing a visuo-motor tracking(3), visual oddball(14), continuous performance(22), and auditory attention tasks(25,27,28).

*Supplementary Table 6. Areas of increased and attenuated activation signal following THC, compared with placebo, during attentional processing*

| *Comparison* | *MNI coordinate* | | | *SDM-Z* | *P* | *Voxels* | *Region* |
|---|---|---|---|---|---|---|---|
| | x | y | z | | | | |
| *THC>PLB* | -4 | 56 | -8 | 3.171 | <0.001 | 1422 | L medial superior frontal gyrus (extending to R medial superior frontal gyrus, L+R gyrus rectus, L+R anterior cingulate/ paracingulate gyri, L+R olfactory cortex) |
| | -52 | 0 | -40 | 2.126 | 0.002 | 34 | L inferior temporal gyrus |
| | 32 | 14 | -14 | 2.001 | 0.004 | 13 | R insula |
| *THC<PLB* | -34 | -2 | 12 | -2.842 | <0.001 | 1556 | L insula (extending to L Rolandic operculum, L superior temporal gyrus, L Heschl gyrus, L lenticular nucleus putamen, L supramarginal gyrus, L postcentral gyrus) |
| | 36 | -4 | 8 | -3.230 | <0.001 | 1266 | R insula (extending to R Rolandic operculum, R lenticular nucleus putamen, R Heschl gyrus, R striatum, R postcentral gyrus, R precentral gyrus, R superior temporal gyrus, R inferior frontal gyrus) |
| | 0 | -60 | -8 | -2.490 | <0.001 | 481 | Cerebellum vermic lobule IV/V |
| | -58 | 0 | 22 | -2.283 | 0.001 | 51 | L precentral gyrus (extending to L postcentral gyrus) |
| | 10 | -76 | 16 | -1.912 | 0.005 | 42 | R calcarine fissure |
| | -8 | -4 | 4 | -2.215 | 0.001 | 38 | L thalamus (extending to L caudate nucleus) |
| | 16 | -26 | 12 | -2.195 | 0.001 | 15 | L thalamus |

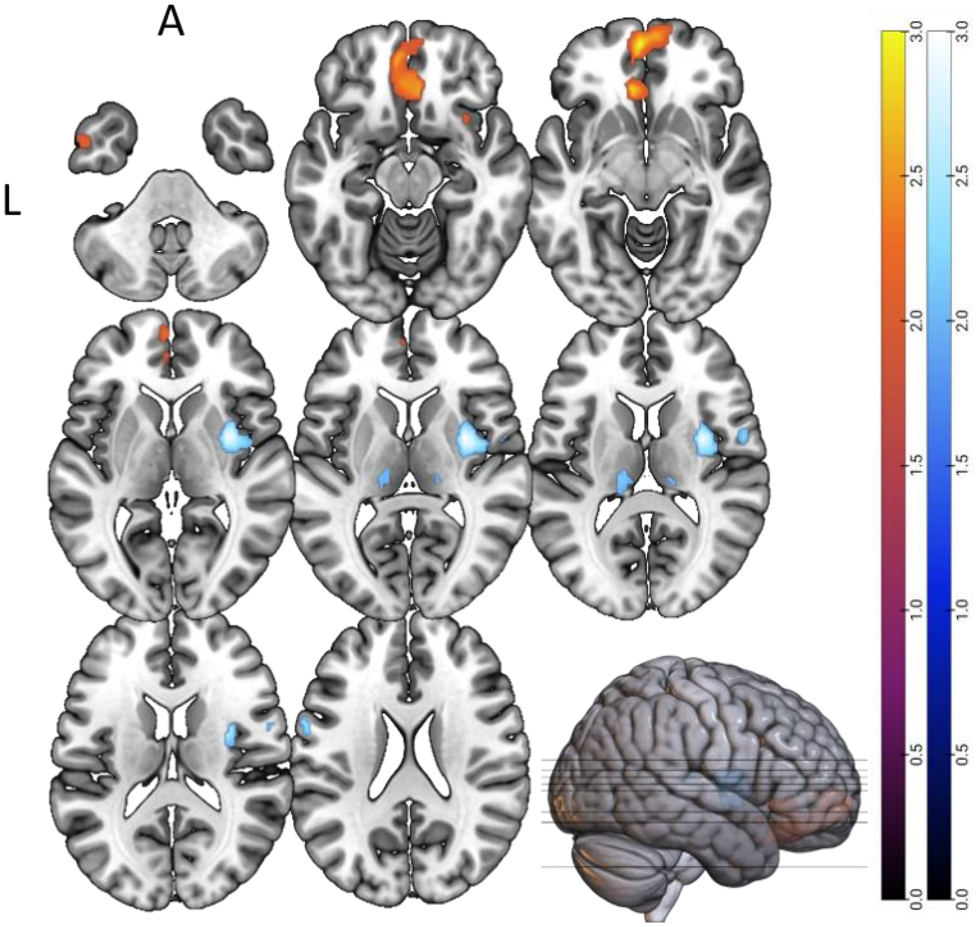


*Supplementary Figure 21. Attention processing: a) Areas of increased activation signal after THC compared with placebo, b) Areas of attenuated activation signal after THC compared with placebo*

*fMRI subgroup*

The fMRI subgroup was composed of 298 participants(3,4,14–19,23,24,29–33).

*Supplementary Table 7. Areas of increased and attenuated activation signal following THC, compared with placebo, during fMRI*

| *Comparison* | *MNI coordinate* | | | *SDM-Z* | *P* | *Voxels* | *Region* |
|---|---|---|---|---|---|---|---|
| | x | y | z | | | | |
| *THC>PLB* | 46 | -76 | 20 | 2.937 | <0.001 | 301 | R middle occipital gyrus (extending to R superior occipital gyrus, R angular gyrus) |
| | 32 | -88 | -8 | 2.549 | <0.001 | 160 | R inferior occipital gyrus (extending to R middle occipital gyrus) |
| | 8 | 62 | -4 | 2.499 | <0.001 | 109 | R superior frontal gyrus, medial orbital (extending to R superior frontal gyrus, orbital part) |
| | -32 | -88 | 20 | 2.305 | <0.001 | 112 | L middle occipital gyrus |
| | -28 | 0 | 56 | 1.949 | 0.002 | 61 | L middle frontal gyrus (extending to L superior frontal gyrus, dorsolateral, L precentral gyrus) |
| | -24 | -22 | 54 | 2.216 | <0.001 | 31 | L precentral gyrus |
| | 12 | 22 | 30 | 2.092 | <0.001 | 23 | R median cingulate / paracingulate gyri (extending to R anterior cingulate gyrus) |
| | 0 | -20 | -12 | 2.1 | 0.001 | 20 | L thalamus |
| *THC<PLB* | -44 | -12 | 8 | -2.357 | 0.001 | 114 | L insula (extending to L rolandic operculum, L Heschl gyrus, L superior temporal gyrus) |
| | -62 | 6 | 24 | -2.343 | 0.001 | 72 | L precentral gyrus (extending to L inferior frontal gyrus, opercular part, L postcentral gyrus) |
| | -52 | -60 | 44 | -2.374 | <0.001 | 70 | L inferior parietal lobule (extending to L angular gyrus) |
| | -42 | 8 | -24 | -2.32 | 0.001 | 21 | L temporal pole, superior temporal gyrus (extending to L temporal pole, middle temporal gyrus) |
| | -64 | -56 | 8 | -2.276 | 0.001 | 17 | L middle temporal gyrus |
| | -36 | 0 | -38 | -2.171 | 0.002 | 15 | L inferior temporal gyrus |
| | 8 | -64 | -52 | -2.079 | 0.003 | 10 | R cerebellum, hemispheric lobule VIII (extending to R lobule IX) |

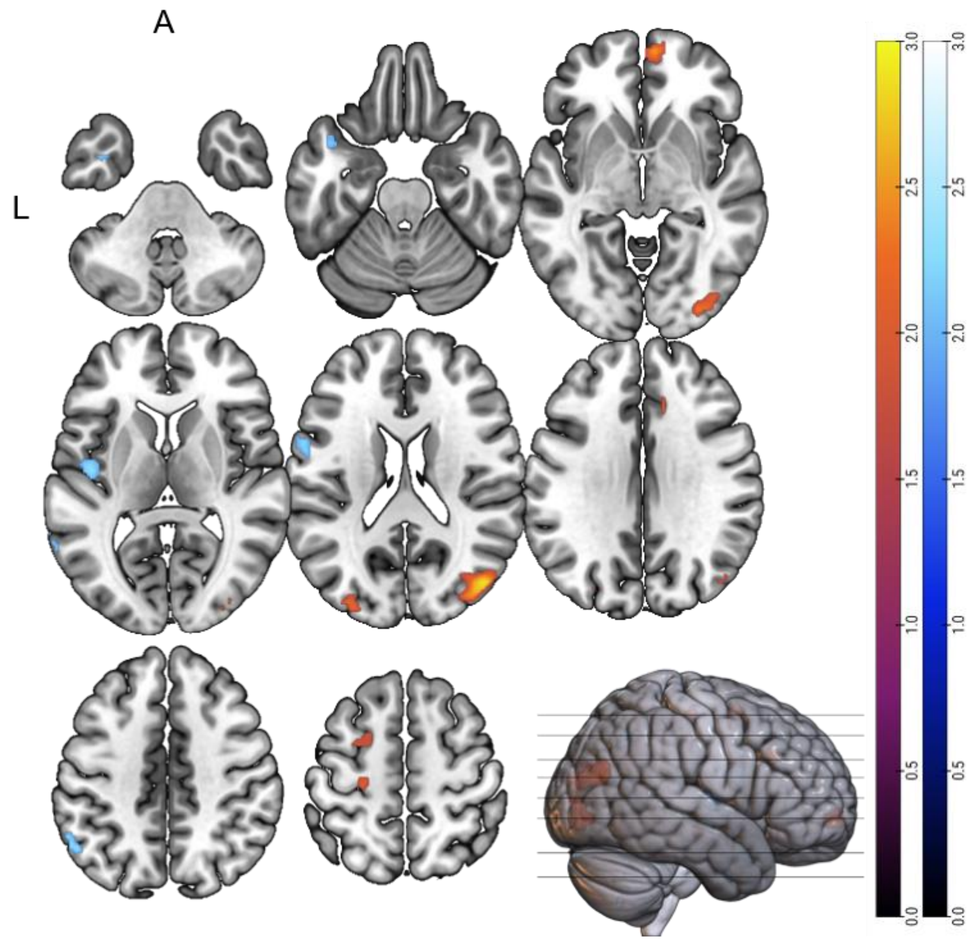


*Supplementary Figure 22. fMRI contrasts: a) Areas of increased activation signal after THC compared with placebo, b) Areas of attenuated activation signal after THC compared with placebo*

#### *THC isolate subgroup*

The THC isolate subgroup was composed of 284 participants(14–24,29–33).

*Supplementary Table 8. Areas of increased and attenuated activation signal following THC isolate, compared with placebo*

| *Comparison* | *MNI coordinate* | | | *SDM-Z* | *P* | *Voxels* | *Region* |
|---|---|---|---|---|---|---|---|
| | x | y | z | | | | |
| *THC>PLB* | 44 | -76 | 20 | 3.073 | <0.01 | 624 | R middle occipital gyrus (extending to R inferior & superior occipital gyrus, R angular gyrus) |
| | -40 | 2 | 28 | 2.338 | <0.01 | 146 | L inferior frontal gyrus, opercular part (extending to L precentral gyrus, middle frontal gyrus) |
| | -32 | -88 | 20 | 2.473 | <0.01 | 105 | L middle occipital gyrus |
| | -42 | -68 | -4 | 2.379 | <0.01 | 79 | L inferior occipital gyrus (L middle occipital gyrus) |
| | 0 | -20 | -12 | 2.319 | <0.01 | 25 | L thalamus |
| | 32 | -26 | -12 | 2.131 | 0.002 | 24 | R hippocampus |
| | 12 | 26 | 32 | 2.175 | 0.002 | 17 | R median cingulate / paracingulate gyri |
| | -12 | -74 | -44 | 2.316 | <0.01 | 17 | L cerebellum, lobule VIIB (extending to lobule VIII) |
| | -24 | -22 | 54 | 2.186 | 0.001 | 13 | L precentral gyrus |
| | -32 | -88 | -8 | 2.114 | 0.002 | 12 | L inferior occipital gyrus (extending to L middle occipital gyrus) |
| | 44 | -36 | 8 | 2.216 | 0.001 | 11 | R superior temporal gyrus |
| *THC<PLB* | -56 | -60 | 42 | -2.286 | <0.01 | 194 | L angular gyrus (extending to inferior parietal, excluding supramarginal, gyri) |
| | -8 | 18 | -12 | -2.394 | <0.01 | 88 | L gyrus rectus (extending to L striatum, L+R olfactory cortex, L caudate nucleus, L anterior cingulate/ paracingulate gyri, L gyrus rectus |
| | 8 | -64 | -52 | -2.231 | <0.01 | 88 | R cerebellum, lobule VIII (extending to R cerebellum lobule IX & VIII) |
| | -36 | 0 | -38 | -2.33 | <0.01 | 85 | L inferior temporal gyrus (extending to L fusiform gyrus, L temporal pole, superior temporal gyrus, L parahippocampal gyrus) |
| | -6 | -54 | 72 | -2.186 | <0.01 | 79 | L precuneus |
| | -64 | -56 | 8 | -2.205 | <0.01 | 60 | L middle temporal gyrus |
| | -44 | 8 | -24 | -2.102 | 0.001 | 20 | L temporal pole, superior temporal gyrus (extending to L temporal pole, middle temporal gyrus) |
| | 68 | -18 | -8 | -2.025 | 0.002 | 18 | R middle temporal gyrus (extending to R superior temporal gyrus) |
| | -4 | 32 | 60 | -2.136 | 0.001 | 11 | L superior frontal gyrus, medial (extending to L supplementary motor area) |

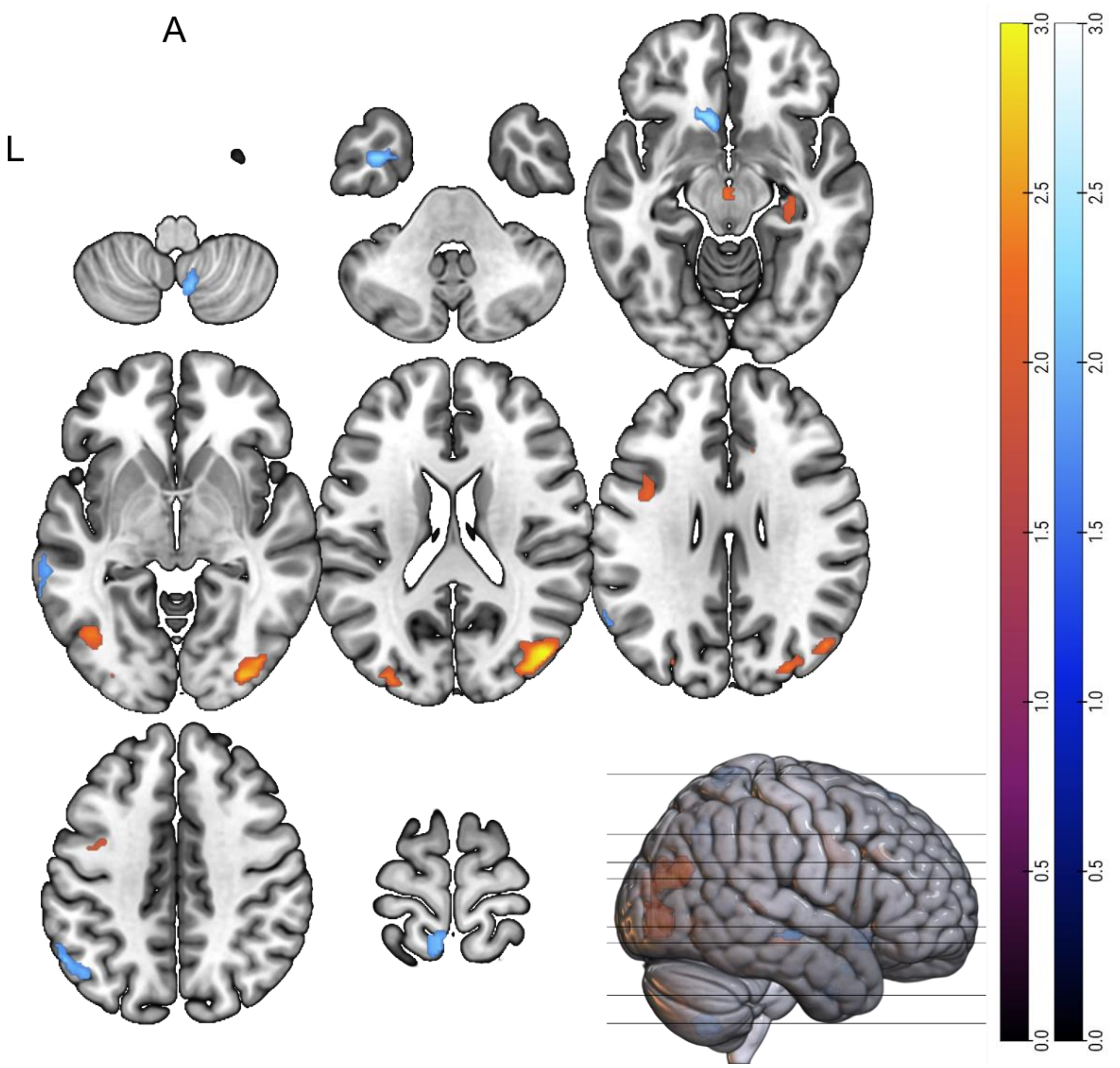


*Supplementary Figure 23. THC isolate: a) Areas of increased activation signal after THC compared with placebo, b) Areas of attenuated activation signal after THC compared with placebo*

### *Supplementary scanner strength comparison analyses*

A comparative analysis was conducted to examine differences in brain activation signal across subjects from those where were scanned in a 3T scanner with those in a 1.5T scanner.

*Supplementary Table 9. Areas of increased and attenuated activation signal following THC, compared with placebo when comparing 3T MRI scanners with 1.5T*

| *Comparison* | *MNI coordinate* | | | *SDM-Z* | *P* | *Voxels* | *Region* |
|---|---|---|---|---|---|---|---|
| | x | y | z | | | | |
| *THC>PLB* | 62 | -32 | 12 | 3.263 | <0.001 | 888 | R superior temporal gyrus (extending to R rolandic operculum, R Heschl gyrus, R supramarginal gyrus, R temporal pole, superior temporal gyrus) |
| | 56 | 18 | 14 | 3.141 | <0.001 | 499 | R inferior frontal gyrus opercular part (extending to R inferior temporal gyrus, triangular part, R precentral gyrus, R rolandic operculum, R precentral gyrus) |
| | -42 | -32 | 12 | 2.978 | <0.001 | 411 | L superior temporal gyrus (extending to L rolandic operculum, L heschl gyrus, L insula) |
| | -30 | -88 | 12 | 2.74 | <0.001 | 355 | L middle occipital gyrus (extending to L inferior occipital gyrus) |
| | -4 | -80 | 12 | 2.759 | <0.001 | 304 | L calcarine fissure (extending to L lingual gyrus) |
| | 44 | -76 | 20 | 2.985 | <0.001 | 241 | R middle occipital gyrus (extending to R middle temporal gyrus) |
| | -40 | 6 | 30 | 2.62 | 0.001 | 243 | L precentral gyrus (extending to L inferior frontal gyrus, opercular & triangular part, L middle frontal gyrus) |
| | -44 | -68 | -4 | 2.782 | <0.001 | 126 | L inferior occipital gyrus (extending to L middle temporal gyrus, L middle occipital gyrus) |
| | 36 | -84 | 2 | 2.6 | 0.001 | 97 | R middle occipital gyrus |
| | 56 | -28 | 40 | 2.595 | 0.001 | 61 | R supramarginal gyrus (extending to R postcentral gyrus) |
| | 36 | -4 | 16 | 2.493 | 0.001 | 53 | R insula (extending to R lenticular nucleus, putamen, R striatum) |
| | -30 | -12 | 4 | 2.591 | 0.001 | 22 | L lenticular nucleus, putamen (extending to L striatum) |
| | -48 | -54 | 0 | 2.41 | 0.002 | 19 | L middle temporal gyrus |
| | -16 | -80 | 32 | 2.354 | 0.003 | 10 | L superior occipital gyrus |
| *THC<PLB* | 4 | 52 | -12 | -2.969 | <0.001 | 2824 | R superior frontal gyrus medial orbital (extending to L+R anterior cingulate/ paracingulate gyri, L+R gyrus rectus, L+R olfactory cortex, , L+R striatum, , L+R medial & medial orbital superior frontal gyrus, R dorsolateral superior frontal gyrus, L caudate nucleus) |
| | -40 | 0 | -38 | -2.819 | <0.001 | 980 | L inferior temporal gyrus (extending to L temporal pole, middle temporal gyrus, L fusiform gyrus, L fusiform gyrus, L parahippocampal gyrus, L temporal pole, superior temporal gyrus) |
| | 52 | 12 | -36 | -2.351 | <0.001 | 369 | R temporal pole, middle temporal gyrus (extending to R inferior & middle temporal gyrus, R temporal pole, superior temporal gyrus) |
| | 24 | 8 | -26 | -2.154 | <0.001 | 241 | R parahippocampal gyrus (extending to R inferior orbital frontal gyrus, R temporal pole |

| | | | | | | |
|---|---|---|---|---|---|---|
| | | | | | | superior temporal gyrus, R insula, R olfactory cortex) |
| -6 | -54 | 72 | -1.864 | <0.001 | 84 | L precuneus |
| 8 | -64 | -50 | -1.896 | <0.001 | 72 | R cerebellum, hemispheric lobule VIII (extending to R lobule IX & VIII) |
| -52 | -60 | 44 | -1.756 | 0.002 | 38 | L inferior parietal lobule (extending to L angular gyrus) |
| -28 | 20 | -24 | -1.89 | 0.001 | 21 | L inferior frontal gyrus, orbital part |
| 12 | -40 | 56 | -1.659 | 0.003 | 14 | R paracentral lobule (extending to R precuneus) |
| 34 | -76 | -36 | -1.685 | 0.002 | 12 | R cerebellum, crus I (extending to crus II) |

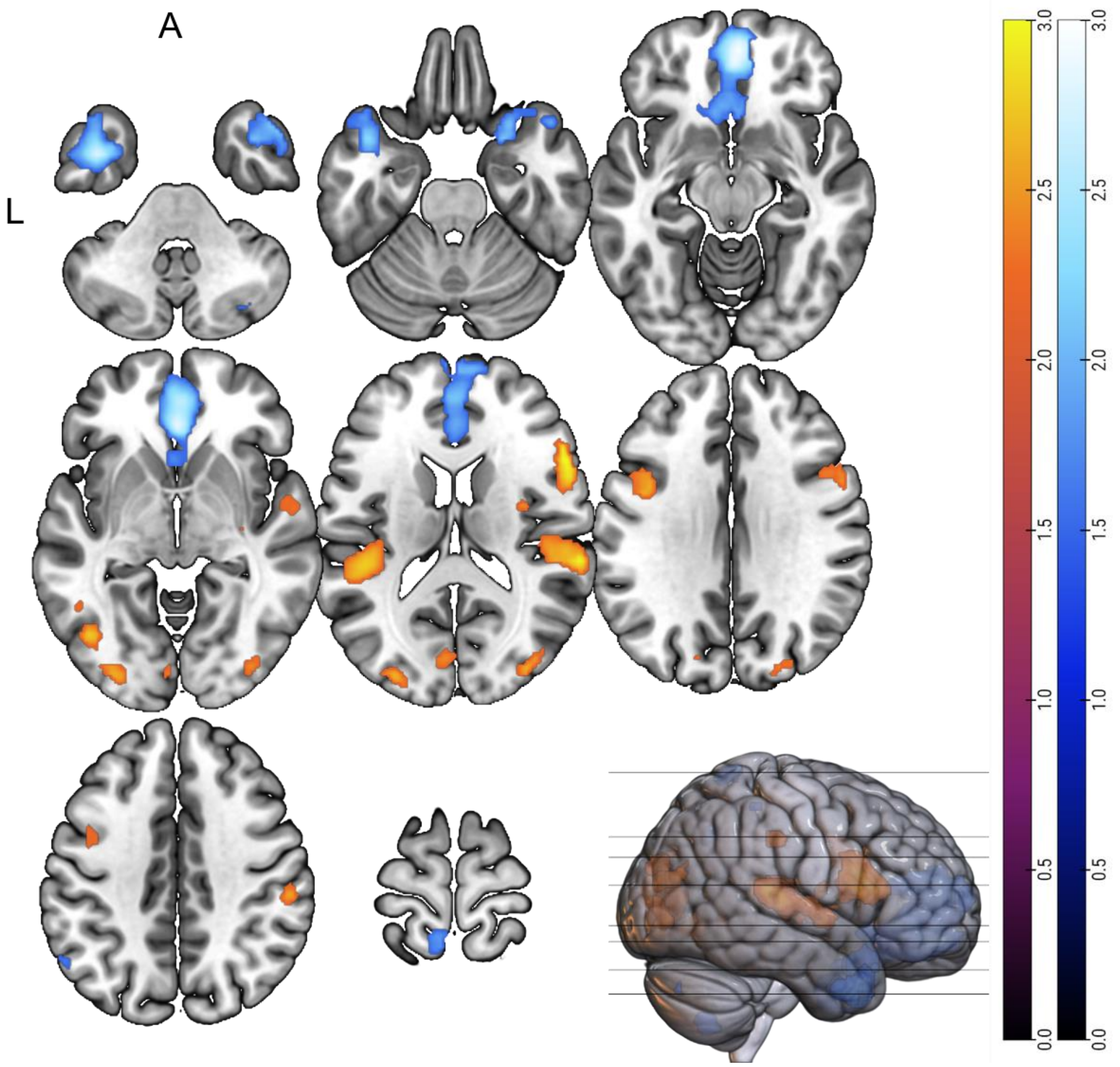


*Supplementary Figure 24. a) increases in brain activation signal across subjects from those where were scanned in a 3T scanner with those in a 1.5T scanner after THC compared with placebo, b) attenuation in brain activation signal across subjects from those where were scanned in a 3T scanner with those in a 1.5T scanner after THC compared with placebo.*

#### *Supplementary main findings with fMRI comparison analyses*

Combining different imaging modalities may be a source of heterogeneity. Therefore, to ensure these factors minimally influenced our core findings, we conducted subgroup analysis of fMRI studies (Supplementary Table 7). We visually inspected the overlap between the findings of our main results and those from the fMRI subgroup alone (Supplementary Figure 25).

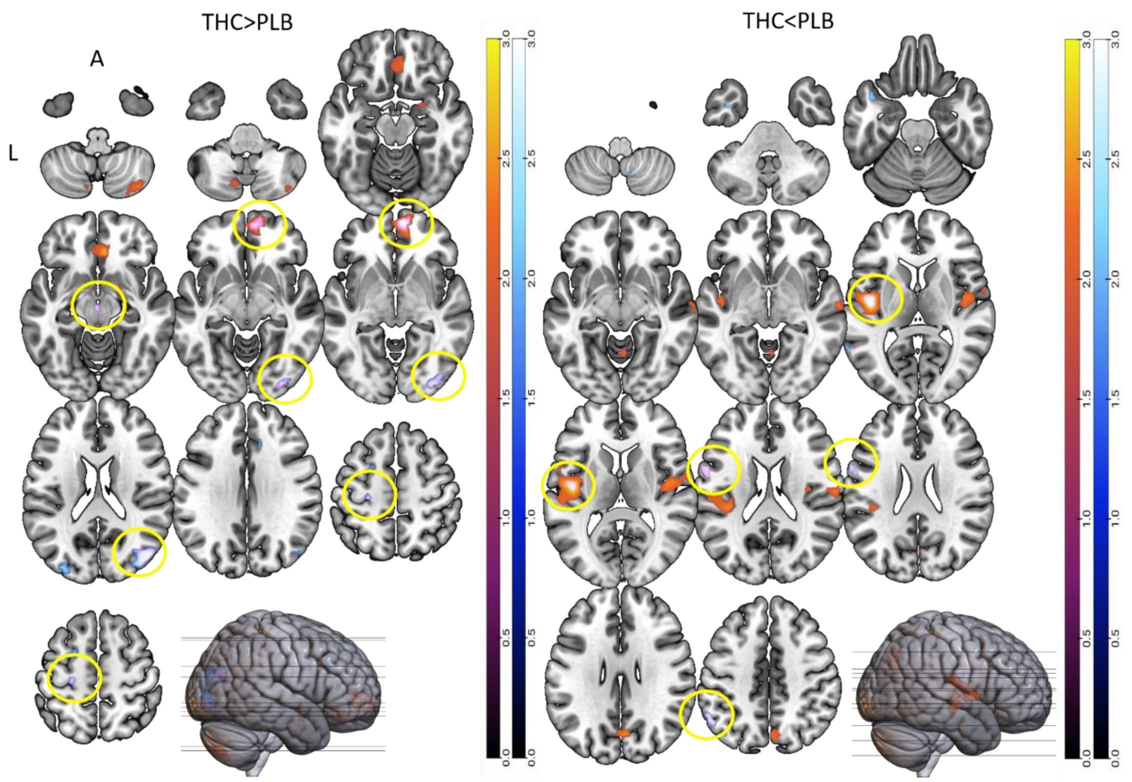


*Supplementary Figure 25. Overlay of the main finding and fMRI maps. Yellow circles highlight clusters of overlap between the maps. Left = brain regions significantly increased by THC. Right = brain regions significantly attenuated by THC. Clusters shown are from both main findings and fMRI subgroup analysis.*

***Supplementary Discussion:***

***Supplementary Discussion 1***

A key aim of the present study was to examine, at the meta-analytic level, to what extent THC has effects on components of a common core network of brain regions, that has been described as a 'domain-general' core network that facilitates cross-task cognitive function(34). Results of the jack-knife sensitivity analyses further seem to be consistent with the idea of a core network of regions modulated by THC as the superior frontal gyrus and the middle temporal gyrus survive all iterations of this leave one study out at a time approach. However, the effects of THC on brain signal that are not part of the core network described by Shine et al., appear to, for the most part, align with the medial frontoparietal and midcingulo-insular (also known as salience) networks(35). These regions include the medial prefrontal/ orbitofrontal cortex, parts of superior and inferior frontal gyri, superior temporal gyrus, hippocampus and precuenus as well as the insula. The medial frontoparietal network has been proposed as responsible for a number of functions including the formation, temporal binding, and dynamic reconfiguration of associative representations based on current goal-states(35), the integration of concepts with brainstem systems to coordinate organism-wide emotional behaviour and generate affective meaning(36) and goal-directed cognition(37,38). The medial frontoparietal network has also been associated with predictive coding, semantic associations, and environmental assessment(39), while the midcingulo-insular network may have a role in the detection of behaviourally relevant stimuli(35). Modulation of these networks may therefore underlie the effects of cannabis use on cognitive processes such as memory(40) and reward processing(41) as well as the induction of transient psychotic symptoms(41).

Here we further examined the effect of THC on the core network of brain regions in more homogenous subgroups of studies employing related cognitive paradigms. It is worth noting that we did not formally examine the extent of spatial overlap of the common core set of brain regions modulated by THC using data from studies observed in more homogeneous subgroups focusing only on resting or activation conditions.  This is mainly because, there are only a limited number of studies that have investigated THC effects under resting or cognitive activation conditions which limit the inferences that can be drawn on the basis of formal hypothesis testing. Nevertheless, a qualitative comparison of the all-inclusive THC effect map with maps from analyses including studies engaging homogeneous sub-groups of studies indicate that the THC effects on certain key brain regions emerge across these analyses including the striatum, thalamus and the anterior cingulate, which have also been identified as being part of the core network described previously(34) as well as  the hippocampus and inferior frontal gyrus within the medial frontoparietal network(35) and the insula within the midcingulo-insular network(35)

When analysing data from emotional paradigms alone THC modulated the function of 14 regions. Within the pre-defined core network, THC augmented the activation signal relative to placebo, in the striatum (including putamen and caudate), superior temporal gyrus, and the thalamus, while in the superior frontal gyrus, middle temporal gyrus, inferior and superior parietal gyrus, middle occipital gyrus, precuneus, angular gyrus, supplementary motor area, and cerebellum there was an attenuation of activation signal under the influence of THC. Within the medial frontoparietal network(35) we found THC augmented the activation signal in the hippocampus and attenuated it in the inferior frontal gyrus. Augmentation of the insula was also found in the midcingulo-insular network.

Across reward paradigms, THC modulated the function of 20 regions. Within the core network THC increased the activation signal, relative to placebo, in the middle and superior frontal and middle temporal gyri, anterior cingulate, striatum, and the cerebellum. In the middle frontal, middle and superior temporal, middle and superior occipital gyri (spatially distinct from increase in brain signal), fusiform gyrus, and inferior parietal lobule there was a decrease of activation signal following THC. No effects of THC on brain signal activation overlapped with the medial frontoparietal network or midcingulo-insular network(35).

During sensory paradigms, THC modulated the function of 6 regions. Within the core network THC augmented the activation signal, compared with placebo, in the middle temporal gyrus, lingual gyrus, and fusiform gyrus while in the caudate nucleus there was a reduction of activation signal under THC. No effects of THC on brain signal activation overlapped with the medial frontoparietal network or midcingulo-insular network(35).

Across memory paradigms, THC modulated the function of 21 regions. Within the core network THC increased the activation signal, relative to placebo, in the middle frontal and superior frontal gyri, middle temporal and occipital gyri, anterior cingulate, and the cerebellum. In the inferior and superior parietal gyri, superior occipital lobule, fusiform and lingual gyri, cuneus, precuneus, thalamus, and cerebellum (spatially distinct from increase in brain signal) there was an attenuation of activation signal under THC. Within the medial frontoparietal network(35) THC effects were found in the inferior frontal gyrus and attenuation of brain signal was observed in the parahippocampus and the hippocampus. Attenuation of the insula was also found in the midcingulo-insular network.

Attentional salience paradigms resulted in the modulation of  the functioning of 10 regions following THC. Within the core network the superior frontal gyrus and the anterior cingulate were augmented following THC administration. Reduction of activation signal was observed within the superior temporal gyrus, striatum, thalamus, and cerebellum post THC. Effects of THC were not observed to overlap with the medial frontoparietal network(35). THC was also found to both augment and attenuate spatially distinct clusters of the insula within the midcingulo-insular network.

***Supplementary Discussion 2***

Our second major finding was that the effect of THC on the pooled effect-size of regional brain signal was related to a proxy measure of regional CB1R density. The first and key evidence for CB1Rs mediating the effects of THC came from a series of blocking studies, where pre-treatment with the CB1R antagonist (inverse agonist) rimonabant blocked the acute psychological and physiological effects of smoked cannabis without altering THC pharmacokinetics(42). Outside of the THC literature, experiments with rimonabant also showed that it can induce emotional memory biases(43) and other alterations to cognition such as reward processesing(44), and has further depressogenic and anxiogenic side effects(45), which has linked CB1R modulation to a diverse array of cognitive and behavioural consequences. However, the only research to date mapping the effects of THC on human brain activation and CB1R availability (as estimated using PET) in the same individuals was conducted by our group, where we showed that the acute effects of THC on anxiety and amygdala function are related to local availability of CB1 receptors(16), although these analyses were spatially restricted to the amygdala. While these converging lines of evidence pointed to the mechanisms underlying the cognitive, behavioural, and neural effects of THC, the current meta-analysis represents the first material demonstration that, across the brain, the neurophysiological effects of THC are spatially correlated with CB1R expression.

***Supplementary Discussion 3***

Our collective findings may also shed light on why heavier cannabis use and higher THC doses are associated with greater deleterious cognitive and behavioural effects(46–48). For example, the fact that THC can have profound effects on memory, attention, psychomotor control, general cognitive function and symptoms not only of psychosis but also depression and anxiety suggests that THC has emotion and cognition-agnostic effects. While such phenomena could never be reduced or attributed to a single brain region, the frontal cortex is well known to orchestrate executive functioning as well as provide top-down control of subcortical structures such as the amygdala, and thus any THC-associated dysfunction here could have wide-ranging effects. The frontal cortex thus provides an apt example: our primary results showed that THC augments activation in a number of frontal regions, and meta-regression analyses further revealed dose-response effects in the same (partially overlapping) regions, such that the higher the THC dose, the greater the effect of THC. While it is already well established that higher doses of THC (or heavier cannabis use) have greater impairing effects on cognition and behaviour, the various threads of our findings together add mechanistic insight into the neurophysiological substrates by which these effects occur.

***Supplementary Discussion 4***

It is important to note that in our analysis Cook’s distance(49) estimate identified the study by Battistella et al.,(3) as being a potential outlier. While outlier/ influence diagnostics are popular in meta-analysis others have recommended against their routine use(50) particularly because of challenges in distinguishing true outliers (where the data is erroneous) from large errors in sampling(51). Instead, they may be seen as a method of sensitivity analysis and inform confidence in findings(52). In the present case, this is especially important because Battistella et al., was an influential study administering the strongest dose of THC (42mg), which may be considered as a large sampling error. Therefore, to consider the results of the Battistella study as outliers would be based on the false assumption that sampling error cannot produce values that extreme. The result would be an underestimation of the variance of the observed values, resulting in underestimation of the population variance. Consequently, valid data points with large sampling errors would have been discarded(51). Hence, we refrained from excluding Battistella et al., from our analyses, but have reported that it may have been a potential outlier and advise appropriate caution in the interpretation of the dose response association results.

***Methodological considerations & heterogeneity***

It may be argued that meta-analytic combination of data from a number of functional neuroimaging studies that use a disparate set of task (activation) or resting state paradigms is not appropriate as that does not allow the detection of task-specific effects but rather more domain-general effects. However, we adopted this approach as the purpose of the present endeavour is to really focus on these domain general effects of THC on brain function that may in turn underlie some of its cognitive domain (task) specific effects. Our results demonstrate that these domain general effects of THC are directly related to a proxy measure for the local availability of its target receptor in the brain, indicating the validity of this analytic approach. We also, however, carried out subgroup analysis focusing on studies that employed similar cognitive tasks. Although, it must be acknowledged that these subsets contained few data sets thus reducing the power of these sub-analyses.

A further limitation of this study is that we used mRNA expression to approximate cannabinoid gene expression. Therefore, we report an index of gene *transcriptional* activity as an indirect proxy of cannabinoid receptors, which is ultimately determined by gene *translation*. This distinction is important as previous reports highlight significant variance between mRNA and protein levels within a tissue(53–55). Moreover, gene expression (transcriptional activity) and protein abundance (translational activity) are not always positively correlated(56,57). This limitation is coupled with the caveat that the spatial of association between CB1R and the effect size estimate may not necessarily imply a causal association.

Furthermore, there are certain limitations inherent to meta-analytic integration of neuroimaging data. In this meta-analysis, we employed an approach that utilised both t-maps and coordinates. The integration of coordinates into our analysis, as opposed to t-maps alone, may have increased the risk of bias in our results as coordinates are most often reported to a threshold of $p<0.05$. This stringent threshold may have excluded clusters, which when pooled with other results from other studies, may have found a significant alteration in activation. Nonetheless, we attempted to mitigate this issue by including as many t-maps as possible.

General sources of heterogeneity consisted of comorbid exposure to alcohol, nicotine and other drugs between participants included in the various studies. Moreover, many studies did not control for the length of cannabis abstinence prior to scan acquisition, an issue when considering THC sensitisation. Broadly, participant cannabis history can be grouped into 1 to 25 uses in their lifetime(3,14–17,29,33) and 30 to 1415 uses in their lifetime(4,18,30,19–23,25,27,28). One study was conducted within cannabis naïve participants(24), one was conducted within those with nicotine addiction(23), and one used half a sample of those with chronic cannabis use(26). This heterogeneity is an accepted limitation, however, to assess the influence of individual studies on the main findings we conducted jack-knife leave-one-out sensitivity analysis, as detailed in Supplementary Table 1. The 22 studies included in this analysis were re-analysed, each time excluding 1 single study at a time to establish whether each cluster remained statistically significant. Following this analysis 87% of all clusters survived indicating that our findings are indeed robust. Moreover, when leaving out the study with cannabis naïve participants, 14 out of the 16 clusters survived. Excluding the nicotine study and chronic cannabis study resulted in 12 out of the 16 clusters surviving. Furthermore, QH statistics were assessed in terms of a chi-square distribution and reported after conversion to standard z values to create a map. This map was overlaid on the final mean map for visual inspection of areas of overlapping significant heterogeneity with areas of thresholded activation or attenuation signal. Visual inspection of overlap of meta-analytic activation maps and heterogeneity maps indicated that no areas in our main analysis were influenced by significant heterogeneity.

Nevertheless, it cannot be ruled out that these variables may have influenced brain substrate engagement following drug administration, introducing the risk of type I and II errors, however the large sample size included in this meta-analysis is likely to have mitigated this issue(58).

Further sources of heterogeneity stemmed from combining fMRI and PET data. The PET studies differed greatly in their study design and statistical analysis. The PET studies(25–28) all employed a ‘subtraction analysis’ approach, whereby they subtracted regional blood flow differences following smoking THC-rich cannabis from a control. Subtractions were followed by across subject averaging of the subtraction images and computation of voxel by voxel t-tests of the CBF changes. Significant regions of activation were calculated on the t-map images, using a technique that corrects for the large number of voxel by voxel t-tests performed, the lack of independence between voxels, and the resolution of the processed PET images(59). All t-map analyses used a t-threshold of 3.61 ($p<0.0005$ uncorrected), and a volume threshold of 100 contiguous voxels. This method of analysis differed from the other studies included in this analysis who conducted direct statistical analyses on the THC vs placebo contrasts using either a t-test or ANOVA. Moreover, O'leary and colleagues did not use a placebo control, rather, they compared a potent THC cannabis with cannabis with the THC removed. The extract of *Cannabis sativa* has been reported to contain over 140 different phytocannabinoids, with THC and cannabidiol being the most abundant(60), thus contaminant effects of other cannabinoids may have influenced the results. When the analysis was limited to studies using THC isolate only, as opposed to THC-rich cannabis, new clusters of activation arose in the THC isolate group. This may be due to the absence of other cannabinoids such as cannabidiol, which have shown an opposing effect to THC on brain activity(61,62) which may counteract the effect of THC. Furthermore, while all studies included in the analysis had a minimum one-week wash out period between drug administration two PET studies administered both THC-rich cannabis and THC-removed cannabis on the same day(25,28). To ensure these factors minimally influenced our core findings, we conducted subgroup analysis of fMRI studies, and those of isolated THC. When comparing the subgroup results with the core analysis, some clusters from the core analysis survived, suggesting stability in the results.

***Supplementary References:***

7. Egger M, Smith GD, Schneider M, Minder C. Bias in meta-analysis detected by a

simple, graphical test. Br Med J [Internet]. 1997 [cited 2020 Nov 16];315(7109):629–34. Available from: https://pubmed.ncbi.nlm.nih.gov/9310563/

14. Bhattacharyya S, Crippa JA, Allen P, Martin-Santos R, Borgwardt S, Fusar-Poli P, et al. Induction of psychosis by Δ9-tetrahydrocannabinol reflects modulation of prefrontal

and striatal function during attentional salience processing. Arch Gen Psychiatry. 2012;69(1):27–36.

28. O'Leary DS, Block RI, Flaum M, Schultz SK, Boles Ponto LL, Watkins GL, et al. Acute

marijuana effects on rCBF and cognition: A PET study. Neuroreport. 2000 Nov 27;11(17):3835–41.

35. Uddin LQ, Yeo BTT, Spreng RN. Towards a Universal Taxonomy of Macro-scale

Functional Human Brain Networks [Internet]. Vol. 32, Brain Topography. Springer; 2019 [cited 2021 May 18]. p. 926–42. Available from: https://doi.org/10.1007/s10548-019-00744-6

41. Gunasekera B, Diederen K, Bhatthacharyya S. Cannabinoids, reward processing, and psychosis. Psychopharmacology (Berl). 2021;

50. Hunter JE, Schmidt FL. Methods of Meta-Analysis: Correcting Error and Bias in Research Findings. 2nd ed. Sage: Thousand Oaks. CA: JSTOR; 2004.

https://www.nature.com/articles/nature10098
